# Restoration of *PKM1* improves functional maturation of human stem-cell derived-β cell by regulating PEP metabolism

**DOI:** 10.1101/2024.11.14.623532

**Authors:** Haopeng Lin, Deqi Chen, Feng Zhang, Xin Liu, Xiaoxiao Xie, Qifei Dong, Jiawei Yan, Jiaxiang Yin, Zirong Bi, Kuo Jiang, Tongran Zhang, Peng Xue, Wei Peng, Lihua Chen, Tao Xu, Yanying Guo, Zonghong Li, Huisheng Liu

## Abstract

Human stem cell-derived β (SC-β) cells still exhibit limited glucose response required for insulin secretion due to glycolytic bottlenecks, yet how these metabolic abnormalities impact glucose response and functional maturation of SC-β cells remains unclear. In this study, we identified a metabolic checkpoint located at PEP accumulation that impeded the functional maturation, which was rescued by restoration of pyruvate kinase 1 (*PKM1*). Glucose-tracing metabolomics in human stem cell-derived islets revealed abnormal glycolytic PEP accumulation at resting condition, resulting in impaired calcium response and insulin secretion upon high glucose or glycolytic metabolite stimulation. Mechanistically, elevated PEP significantly raised intracellular basal calcium levels, leading to downregulated expression of genes involved in TCA cycle elucidated by single cell transcriptomics. Furthermore, the activity of pyruvate kinase, which metabolizes PEP, was notably reduced due to low PKM1 expression. By overexpressing PKM1, the impairment of TCA-related genes caused by PEP accumulation was reversed via modulating PEP metabolism, resulting in enhanced calcium responses and insulin secretion upon high glucose stimulation. Together, we discovered a novel role of PKM1-regulated PEP metabolism in mediating the functional maturation of human SC-β cells. This study highlights the importance of metabolic reprogramming in human SC-β cell maturation, advancing cell therapy approaches for diabetes treatment.

## Introduction

Functionally mature stem cell-derived β (SC-β) cell is important to ensure the effectiveness of SC-β cell transplantation therapy for diabetic treatment. One of the most important characteristics of pancreatic β functional maturation is glucose-stimulated insulin secretion (GSIS). While recent protocols have shown improvements in GSIS in SC-β cells, they still exhibit a significantly reduced GSIS compared to native human β cells, presenting a barrier to the effectiveness of β cell replacement therapies (*1*, *2*).

In native β cells, glucose metabolism generates ATP through glycolysis and oxidative phosphorylation (OXPHOS), which leads to the closure of K_ATP_ channels, Ca² influx, and subsequent insulin release. (*3*). While SC-β cells can secrete insulin in response to TCA cycle metabolites and depolarization, they are less capable of responding to high glucose levels, possibly due to a metabolic defect located at glycolytic GAPDH and PGK1 protein (*1*, *4*, *5*). Additionally, inhibition of GAPDH leads to the accumulation of upstream metabolites and impair the oxidative phosphorylation necessary for insulin secretion in diabetic β-cells via mTOR1 signaling remodeling (*6*), a pathway also involved SC-β cell maturation (*7*). However, attempts to enhance GAPDH and PGK1 expression in SC-β cells have not restored GSIS (*4*), indicative of unidentified downstream metabolic bottleneck that might decouple the glycolysis-OXPHOS.

Impaired PEP metabolism is another major metabolic obstacle hampering SC-β from proper glucose response (*1*, *4*). Recently, PEP from glycolysis gains more attention for its ability to generate ATP by pyruvate kinase (PK) to initiate K_ATP_ closure, Ca^2+^ influx and triggering pathway via pyruvate kinase, challenging the canonical model of glucose-regulated K_ATP_ channels (*8–10*). Albeit this is still controversial (*11–14*), abnormal PEP metabolism would inevitably impair downstream metabolism required for insulin release (*15*). In SC-β cells, bypassing the PEP cycle with extracellular PEP, oxaloacetic acid (OAA) and other permeable metabolites indeed significantly increased insulin secretion (*4*, *16*). However, the role of PEP metabolism in SC-β function, cell identity and maturation remain unclear.

PKM1 and PKM2 are two main splicing isoforms of pyruvate kinase muscle (PKM) isozymes responsible for converting PEP to pyruvate in pancreatic β cells. While PKM2 activity— regulated by fructose 1,6-bisphosphate (FBP)—accounts for only 10% of PK activity in mature β cells, the constitutively active PKM1 accounts for 90%. Both PK isoforms are essential for PEP metabolism and insulin secretion (*10*). Due to their sequence similarity, most transcriptomic and proteomic studies cannot distinguish between PKM1 and PKM2, leaving it unclear whether a deficiency in either isoform or PK activity could affect glucose metabolism and functional maturation via regulating PEP in SC-β cells.

In this study, we explore the metabolic bottleneck that disconnects the glucose response and its effect on SC-β functional maturation. We identified abnormal PEP accumulation at resting glucose levels as a key checkpoint in SC-β cell metabolism. This accumulation elevated the basal calcium levels independently of K_ATP_ channels, potentially disrupting glucose response and gene expression. Restoration of PKM1, which was barely detectable in SC-β cells, improved GSIS, enhanced the Ca² response, and normalized expression of key glycolytic and OXPHOS genes that had been affected by PEP. These findings reveal new insights into abnormal PEP metabolism and its metabolic programming role in SC-β cell functional maturation.

## Results

### Metabolomic profiling of SC-islets

We first generated SC-islets following established protocols **(Supplementary Figure 1A)** (*2*, *17*, *18*). In agreement with previous studies, SC-islets differentiated from human embryonic stem cell lines H1 and H9 showed increased β-cell specific gene expression (*INS, NKX6.1, MAFA)* and relatively uniform size **(Supplementary Figure 1B, C).** However, SC-islets displayed limited glucose-stimulated insulin secretion (GSIS) compared to mouse islets **(Supplementary Figure 1D).**

To investigate glucose metabolic defect in SC-islets, we compared glucose metabolic fate between mouse- and SC-islets by performing metabolite tracing experiment with labeled [U-¹³C]- glucose **(Figure 1A)**. Key glycolytic intermediates—glucose 6-phosphate (G6P), fructose 6-phosphate (F6P), dihydroxyacetone phosphate (DHAP), and 3-phosphoglycerate (3-PG)/2-phosphoglycerate (2-PG)—showed similar glucose-dependent increases in both mouse and SC-islets, indicating that glucose metabolism from glucose through to the 3-PG/2-PG stage remains relatively intact. **(Figure 1B-E).** Interestingly, while high glucose-induced PEP levels were comparable between mouse and SC-islets, basal PEP levels at low glucose were significantly higher in SC-islets with lower stimulating index, as were pyruvate levels **(Figure 1F-G).** Similarly, the glucose-dependent increase in metabolites from the TCA cycle, such as citrate, α-ketoglutarate (α-KG), malate, ATP and GSH, was also absent in SC-islets **(Figure 1H)**, suggesting that the disrupted glucose metabolism may originate from dysregulated glycolytic PEP metabolism **(Figure 1I)**.

**Figure 1.**
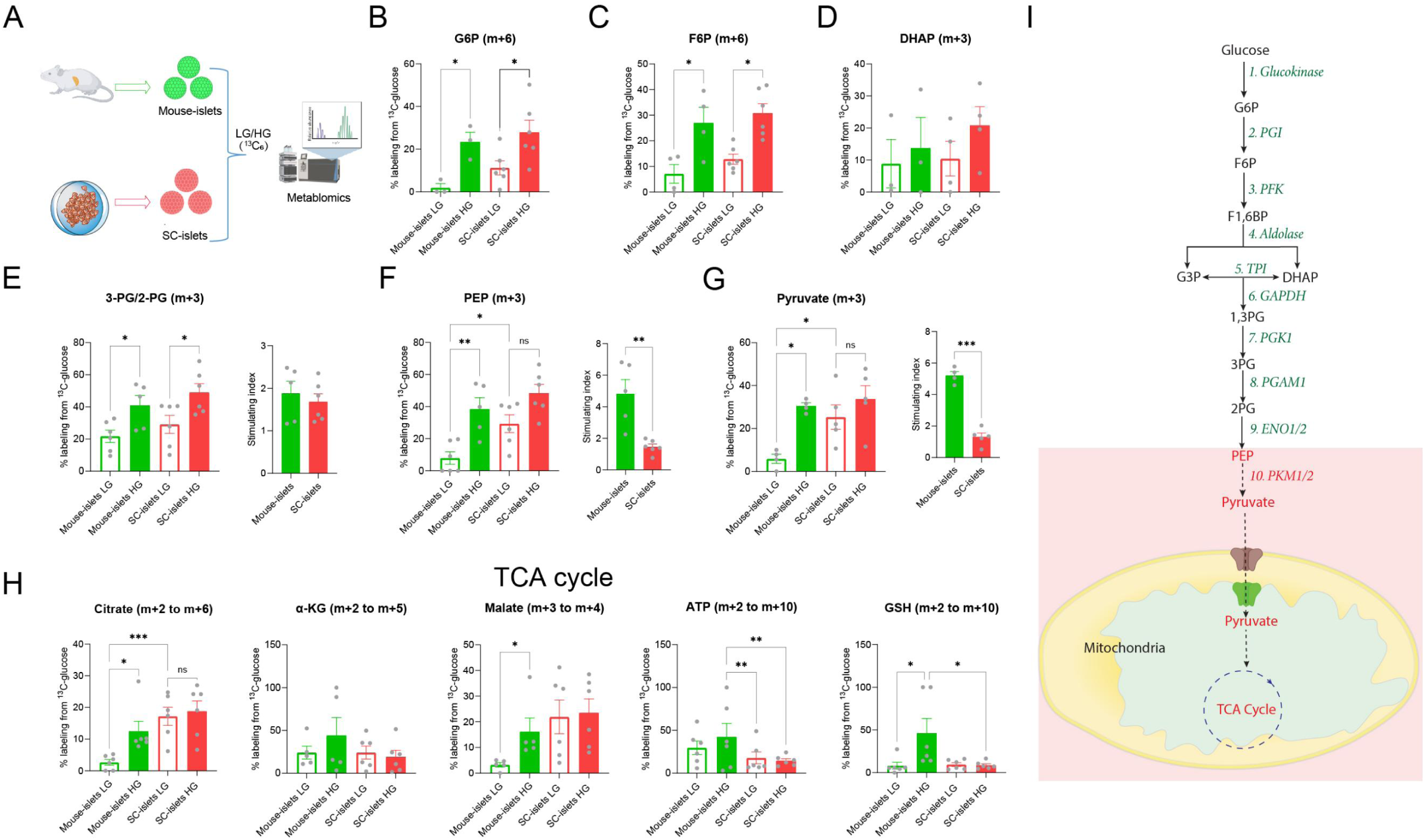
Metabolic tracing analyses of glycolytic and TCA cycle metabolites in mouse- and SC-islets. **(A)** Graphic overview of glucose tracing experimental procedure. Primary mouse islets and SC-islets were labelled with ^13^C at 2.8 mM (LG, low glucose) or 16.7 mM (HG, high glucose) [U^13^C_6_]-glucose for 60 min before LC-MS detection for metabolomics. **(B-G)** Labeled metabolites percentages (percentage of ^13^C_6_ labeling of total metabolites) of glycolytic metabolites. G6P, glucose-6-phosphate; F6P, fructose-6-phosphate; DHAP, dihydroxyacetone phosphate; 3-PG/2-PG, 3-phosphoglycerate/2-phosphoglycerate; PEP, phosphoenolpyruvate. **(H)** Labeled metabolites percentages of TCA cycle metabolites. α-KG, α-Ketoglutarate. (M + x) represents the count of carbon atoms that are labeled with ^13^C. (**I):** Overview of glucose metabolism with metabolic defect starting from PEP in SC-islets compared to mouse islets. All data are presented as mean ± SEM, with statistical significance determined using two-way ANOVA followed by Sidak’s multiple comparison test. ns, not significant. *P < 0.05, **P < 0.01, ***P < 0.001. SC-islets (n = 3–6) and mouse islets (n = 3–4) for experiments.

### Dysregulated PEP metabolism limited glucose-dependent function

To examine if the dysregulated PEP metabolism was the glycolytic bottleneck to prevent glucose response, we measured the effects of PEP and upstream metabolites on SC-β cells. Surprisingly, SC-β cells, differentiated from the H1-Ins-jGCaMP7f cell line for the purpose of specific labelling SC-β cells for calcium imaging (**Supplementary Figure 2A**, Liu et al., manuscript in submission to Stem Cell Research), showed a strong calcium response and insulin secretion in response to PEP, but weaker than mouse-β cells did **(Figure 2A-B)**. In contrast, cell-permeable metabolites glyceraldehyde (GLA, glyceraldehyde 3-phosphate precursor) and dihydroxyacetone (DHA, DHAP precursor), upstream of PEP, elicited minimal changes in calcium and insulin response in SC-β cells **(Figure 2C-F)**, as were glyceric acid (non-phosphorylated precursor to 3-PG and 2-PG) **(Supplementary Figure 2B**-C), indicative of a metabolic clog downstream of 2-PG.

**Figure 2.**
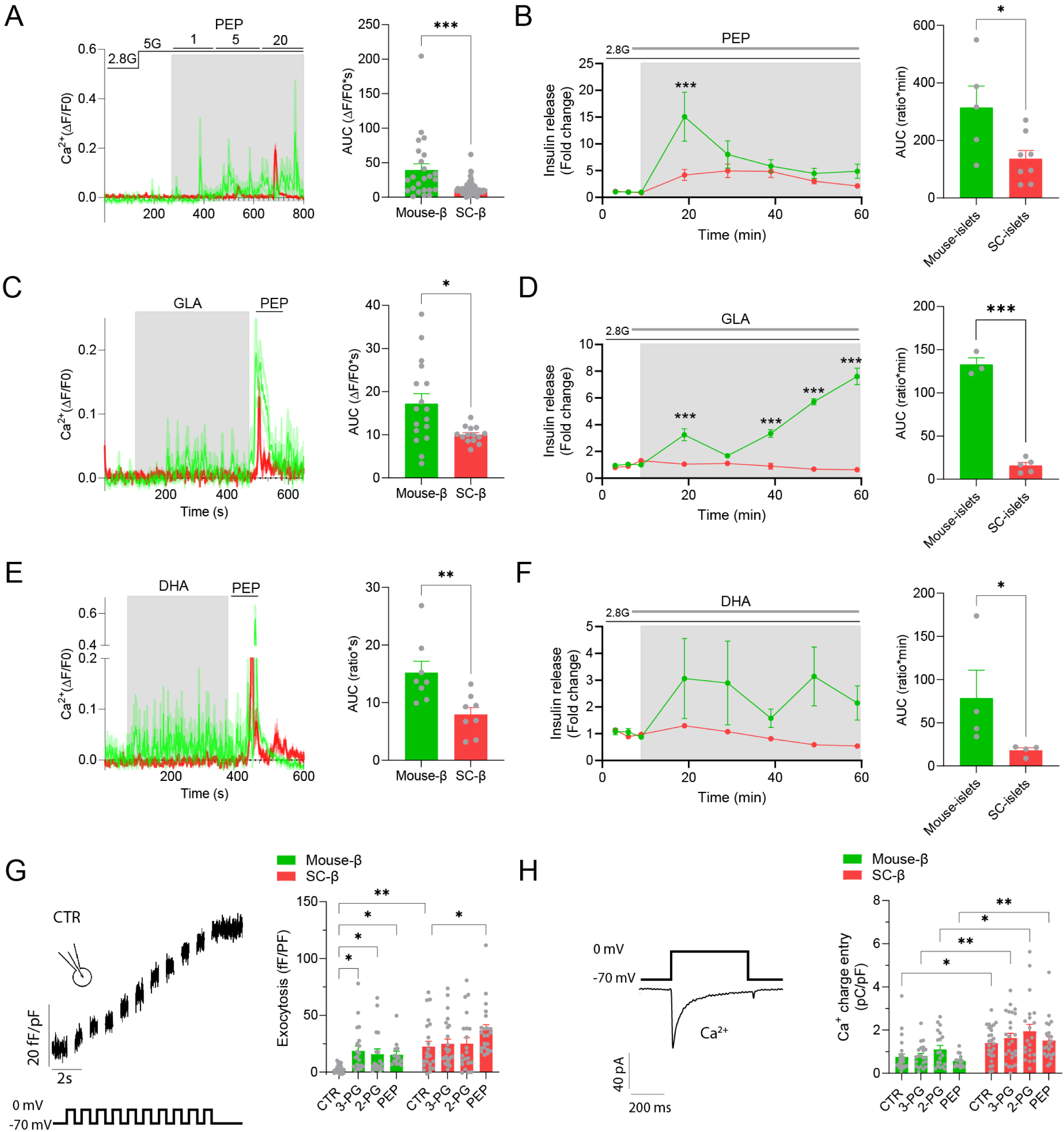
Metabolites upstream of PEP fails to drive functional response in SC-β cells. (**A)** [Ca^2+^]_i_ recordings to different concentration of PEP (1, 5, 10 mM) at 5 mM glucose (5G) from mouse-β (n=24 cells) and SC-β (n=67 cells). (**B)** Insulin secretion to 20 mM PEP at 2.8 mM glucose (2.8G) from mouse-islets (n=5) and SC-islets (n=8). (**C)** [Ca^2+^]_i_ recordings to 20 mM glyceraldehyde (GLA) followed by 20 mM PEP stimulation from mouse-β (n=17 cells) and SC-β (n=14 cells). **(D)** Insulin secretion to 20 mM GLA from mouse-islets (n=3) and SC-islets (n=5). **(E)** [Ca^2+^]_i_ recordings to 20 mM dihydroxyacetone (DHA) followed by 20 mM PEP stimulation from mouse-β (n=8 cells) and SC-β (n=8 cells). **(D)** Insulin secretion to 20 mM DHA from mouse-islets (n=4) and SC-islets (n=4). **(G)** Representative traces (left), and average total normalized exocytotic responses of β-cell (right) with infusion of 5 mM pH-corrected 3-PG, 2-PG and PEP from mouse-β (n=26, 21, 18, 14 cells) and SC-β (n=22, 24, 21, 22 cells) at 2.8 mM glucose. **(H)** Representative traces (left), and average Ca^2+^ currents of β-cell (right) with infusion of 5 mM pH-corrected 3-PG, 2-PG and PEP from mouse-β (n=27, 22, 21, 16 cells) and SC-β (n=26, 25, 20, 22 cells). Glucose level was 2.8 mM unless indicated. For [Ca^2+^]_i_ recordings, single mouse-β cell was loaded with Cal-500AM and identified by immunostaining for insulin. Single SC-β cell was identified by using cells differentiated from H1-Ins-jGCaMP7f. [Ca^2+^]_i_ recordings were analyzed using smooth dynamics method (*46*). For patch clamp, single SC-β cell was identified by using cells differentiated from H1-Ins-EGFP. All data are presented as mean ± SEM, with statistical significance determined using unpaired Student’s t test or two-way ANOVA followed by Sidak’s multiple comparison test. *P < 0.05, **P < 0.01, ***P < 0.001.

Consistently, although 3PG was reportedly not permeable to induce a substantial calcium response in mouse-β cells **(Supplementary Figure 2D)** (*4*), the infusion of 3-PG, 2-PG, or PEP via whole-cell patch clamp significantly enhanced depolarization-induced exocytosis by approximately 9-fold compared to the control group in mouse-β cells. In contrast, SC-β cells, generated from CRISPR-Cas9 method and identified by EGFP **(Supplementary Figure 2E**-G), only showed a very modest PEP-dependent increase (1.5-fold change) in exocytosis compared to the control group **(Figure 2G)**. Remarkably, this was because under control conditions, SC-β cells already exhibited a 10-fold higher exocytosis compared to mouse β-cells, preventing further amplifying effect by the metabolites **(Figure 2G)**. Additionally, SC-β cells had higher amount of depolarization-induced calcium entry across different conditions compared to mouse-β cells **(Figure 2H)**. These data suggested that the saturated effect of accumulated PEP at basal glucose level prevented the glucose- and upstream metabolites-dependent exocytosis.

### PEP elevates Ca²□ levels beyond mitochondrial metabolism and hinders SC-β cell maturation

Since PEP triggered dramatic changes in Ca²^+^ response and insulin secretion, we next explored the underlying mechanism and PEP effect on SC-β cell maturation. We first hypothesized that the effect of PEP was mainly via mitochondrial metabolism. PEP, permeable methyl-pyruvate (MP), and oxaloacetic acid (OAA) at low glucose significantly enhanced insulin secretion to levels higher than those observed with high glucose alone, suggesting insufficient glycolysis upstream of PEP as previously reported (*4*, *16*) **(Figure 3A)**. Surprisingly, while PEP dramatically triggered a dramatic calcium response in both mouse and SC-islets **(Figure 3B)**, MP elicited a calcium response only in mouse islets **(Figure 3C)**. Consistently, PEP triggered a calcium response well beyond those induced by MP and OAA in H1-Ins-jGCaMP7f cell-derived SC-β cells **(Supplementary Figure 2H**-I), indicating that PEP enhanced insulin secretion through a mechanism independent of mitochondrial metabolism. To further investigate this mechanism, we depleted extracellular calcium, and PEP failed to trigger a calcium response **(Figure 3D)**. Previous studies reported that PEP closed K_ATP_ channels by generating ATP via pyruvate kinase (PK), thereby triggering calcium entry (*9*). However, K_ATP_ channel opener diazoxide (Dia) did not dampen the PEP-mediated calcium response **(Figure 3E)**. Additionally, PEP enhanced membrane potential, which remained unaffected by either Dia or pyruvate kinase inhibitor (PKi) **(Figure 3F-G)**. Furthermore, PEP upregulated calcium response and membrane potential in a K_ATP_ channel-free HEK293T cell line **(Figure 2H-I, Supplementary Figure 2J)**, which expressed endogenous L-type voltage-gated calcium channel (*19*). These data suggested that PEP alone dramatically elevated calcium levels even under basal glucose conditions through a K_ATP_-independent pathway, distinct from the mitochondrial pathways used by pyruvate and other TCA metabolites.

**Figure 3.**
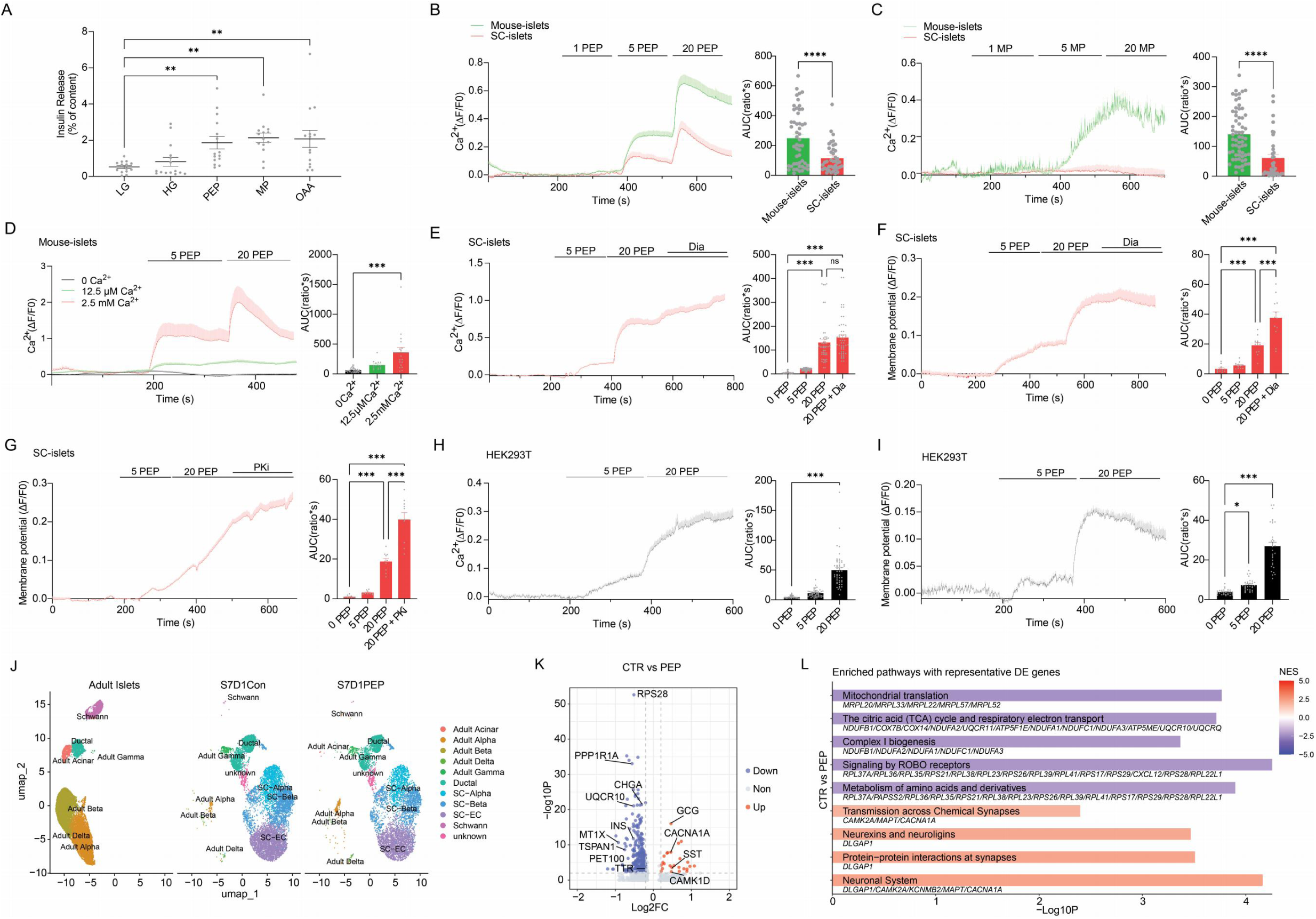
PEP raises calcium response and hinders β cell functional maturation. **(A)** Static insulin secretion to 2 mM glucose (LG), 20 mM glucose (HG), 20 mM PEP, 20 mM MP and 20 mM OAA (n=5, 5 independent experiments in triplicate). **(B)** [Ca^2+^]_i_ recordings to different concentration of PEP (1, 5, 10 mM) at 5 mM glucose (5G) from mouse-islets (n=51 regions from 3 different experiments) and SC- islets (n=40 islet regions from 3 different experiments). **(C)** [Ca^2+^]_i_ recordings to different concentration of MP (1, 5, 10 mM methyl pyruvate) at 5 mM glucose (5G) from mouse-islets (n=63 islet regions from 3 different experiments) and SC- islets (n=35 islet regions from 3 different experiments). **(D)** [Ca^2+^]_i_ recordings at different extracellular concentration of Ca^2+^ (0, 12.5 µM, 2.5 mM) in response to PEP (5, 20 mM) from mouse-islets (n=38, 15, 17 islet regions from 3 different experiments). **(E)** [Ca^2+^]_i_ recordings in response to PEP (5, 20 mM) and diazoxide (Dia, 200 µM) from SC-islets (n=52 islet regions from 3 different experiments). **(F)** Membrane potential recordings in response to PEP (5, 20 mM) and diazoxide (Dia, 200 µM) from SC- islets (n=11 islet regions from 3 different experiments). **(G)** Membrane potential recordings in response to PEP (5, 20 mM) and pyruvate kinase inhibitor (PKi, 10 µM) from SC- islets (n=10 islet regions from 3 different experiments). **(H)** [Ca^2+^]_i_ recordings in response to PEP (5, 20 mM) from HEK293T cells (n=51 cells from 3 different experiments). **(I)** Membrane potential recordings in response to response to PEP (5, 20 mM) from HEK293T cells (n=30 cells from 3 different experiments). [Ca^2+^]_i_ recordings and membrane potential recordings from islets or cells loaded with Fluo-8AM and FluoVolt respectively were analyzed using average fluorescence method (*46*). Recordings were normalized to control condition. Glucose level was 5 mM unless indicated. **(J)** Single-cell RNA-seq transcriptomic profiling of SC-islets cultured with 5 mM PEP throughout stage 6 and collected on day 1 of stage 7 (S7D1). A UMAP-based embedding projection of an integrated dataset comprising 19,435 SC-derived cells and adult human islet cells, separated by sample origin and clustered by cell type. **(K)** Differentially expressed (DE) genes between control (CTR) and PEP-treated β cell group, including adult and SC-β cells. **(L)** Gene sets enriched analysis (GSEA) using Reactome pathways showed upregulated and downregulated pathways along with DE genes between control (CTR) and PEP-treated β cell group. NES, normalized enrichment score. DE, differentially expressed. All data are presented as mean ± SEM, with statistical significance determined using unpaired Student’s t test or one-way ANOVA followed by Sidak’s multiple comparison test. *P < 0.05, **P < 0.01, ***P < 0.001.

Due to the uniqueness of PEP in calcium response, we further investigated the effect of PEP on SC-β cells differentiation SC-islets were cultured with 5 mM PEP throughout Stage 6 and collected for single-cell RNA sequencing at S7D1. We integrated datasets from S7D1-Con, S7D1-PEP, and human adult islets, resulting in a combined dataset consisting of 36,017 cells, including 19,435 cells from human adult islets (*20*). The dataset was clustered based on cell identity, revealing distinct transcriptional profiles for SC-β cells compared to adult islet β cells **(Figure 3J)**. We then focused on SC-β cell subpopulations to investigate the transcriptional changes associated with the PEP effect.

Treatment with PEP disrupted the expression of β-cell-specific genes, such as *CHGA* and *INS*, and also impaired genes related to insulin secretion and exocytosis, including *MT1X* (*21*, *22*), *TSPAN1* (*23*), and *PPP1R1A* (*24*). At the same time, it enhanced the expression of non-β-cell genes, such as *GCG* and *SST* **(Figure 3K, Supplementary Data 2)**. Consistent with these findings, gene set enrichment analysis (GSEA) revealed downregulation of the TCA cycle, the respiratory transport chain, and mitochondrial complex I biogenesis, all of which were necessary for OXPHOS-induced insulin secretion (*25*). ROBO signaling, which played a crucial role in the morphological development, cell identity, and insulin secretion of pancreatic islets, was also downregulated following PEP treatment (*26–28*). Interestingly, PEP treatment induced a neuronal gene system involving calcium signaling (CAMK2A, CACNA1A), a characteristic of immature β-cells, and the removal of this signature was essential for their maturation **(Figure 3L, Supplementary Data 2)** (*29*). Overall, PEP alone induced significantly increased calcium response and insulin secretion, while causing transcriptional changes that hindered further SC-β cell maturation.

### Restoration of *PKM1* improves SC-**β** cell functional response to glucose

As the increase in PEP could reflect either increased production or reduced consumption, we measured the expression and activity of the glycolytic enzymes that might underlie dysregulated PEP metabolism. *ALDOA, PGK1 and PGAM1* showed a trend of reduction in SC-islets, while enolase (*ENO1* and *ENO2*), which directly produced PEP from upstream metabolites, were comparable between mouse- and SC-islet. Remarkably, qPCR data revealed that *PKM1,* which metabolized PEP to pyruvate, was dramatically lower in SC-islets with a difference of approximately 100-fold compared to mouse islets **(Figure 4A)**. Western blotting revealed reduced levels of both inactive (monomer) and active (tetramer) *PKM1* in SC-islets **(Figure 4B)**. Interestingly, *PKM2* expression was higher in SC-islets **(Figure 4A)**, but predominantly in its inactive form (dimer/monomer) compared to mouse islets **(Figure 4B)**. Immunostaining indicated that mouse β cells expressed both PKM1 and PKM2 **(Figure 4C)**, whereas SC-β cells only expressed PKM2 **(Figure 4D)**. Accordingly, pyruvate kinase activity, rather than PGK1 and GAPDH, was significantly lower in SC-islets compared to mouse islets but improved with overexpression of *PKM1* **(Figure 4E, 4F)**. These findings suggest that deficient pyruvate kinase activity, due to insufficient *PKM1* expression and inactive PKM2, likely led to reduced PEP consumption and accumulation.

**Figure 4.**
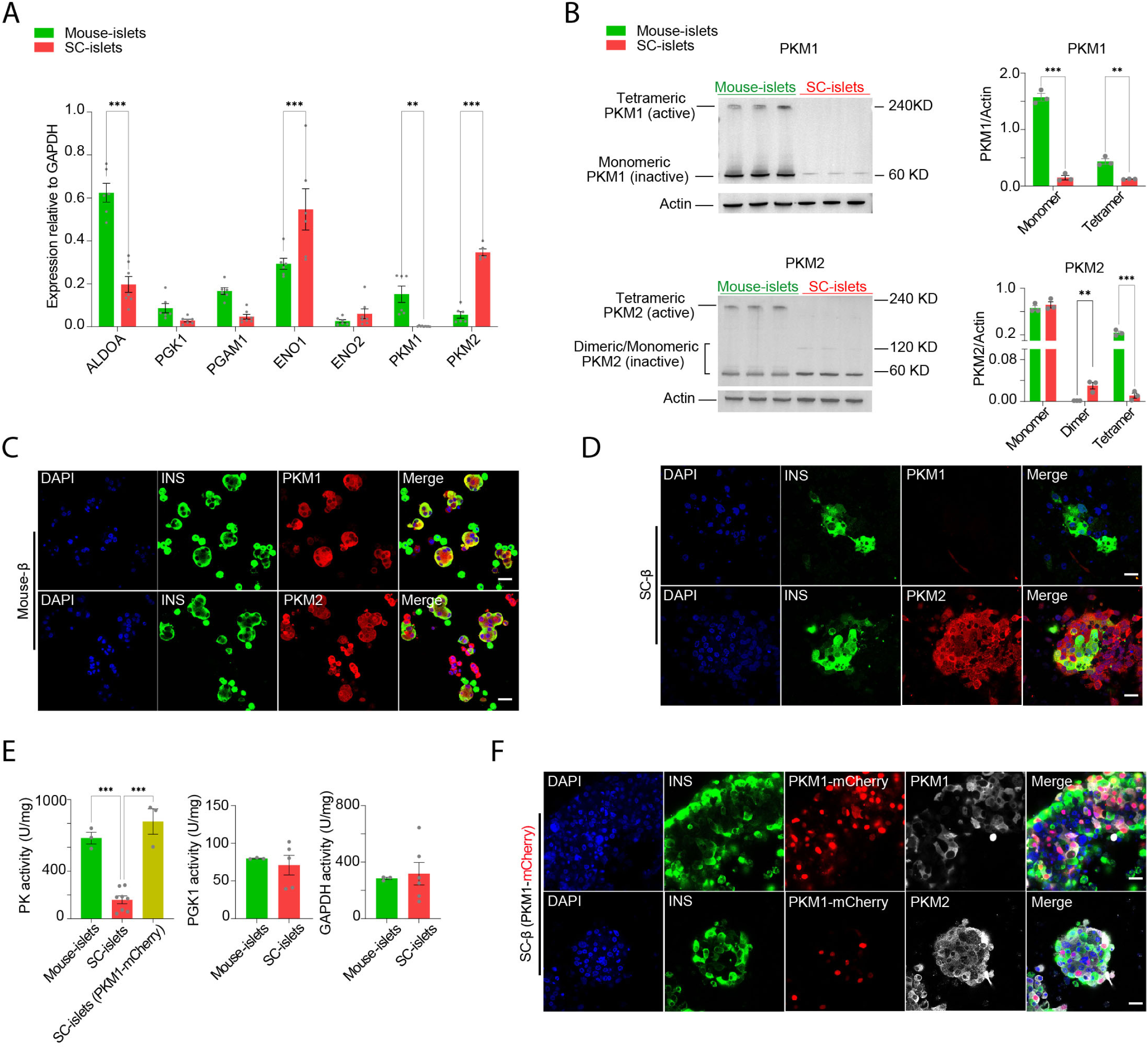
Loss of *PKM1* expression and pyruvate kinase activity in SC-β cells. **(A)** Glycolytic gene expression of mouse- and SC-islets from S7D14 (n=3 different experiments in duplicate). **(B)** Western blotting of PKM1 with normalized band intensity calculation (top) and PKM2 with normalized band intensity calculation (bottom) of mouse- and SC-islets from S7D14 (n=3). **(C)** Representative image of dissociated mouse islet cells immunostained for insulin, PKM1 and PKM2. Scale bar=20 µm. **(D)** Representative image of dissociated SC-islet cells immunostained for insulin, PKM1 and PKM2. Scale bar=20 µm. **(E)** Enzymatic activity of pyruvate kinase (n=3, 8, 3), PGK1 (n=3, 5) and GAPDH (3, 6) of mouse- and SC-islets or overexpressed with *PKM1*-mcherry. **(F)** Representative image of dissociated SC-islet cells immunostained for insulin, PKM1 and PKM2 after *PKM1*-mcherry overexpression. Scale bar=20 µm. All data are presented as mean ± SEM, with statistical significance determined using one-way or two-way ANOVA followed by Sidak’s multiple comparison test. *P < 0.05, **P < 0.01, ***P < 0.001.

We next investigated whether *PKM1* overexpression was sufficient to improve the function of SC-islets. *PKM1*-overexpressing SC-islets showed increased insulin secretion to various glucose levels and FBP, accompanied by heightened calcium responses **(Figure 5A-C)**. On the contrary, acute treatment of PKM2 agonist TEPP-46 (PKa), which significantly boosted calcium responses in mouse β-cells **(Figure 5D)**, failed to enhance calcium or insulin responses in both control and PKM1-overexpressing SC-islets **(Figure 5D-E)**. This indicates that the FBP-induced calcium and insulin increases in PKM1-overexpressing SC-islets were due to metabolic effects of FBP, rather than its effect on PKM2 activation (*30*). We also observed that PKM1 overexpression restored glucose-dependent exocytosis **(Figure 5F)** with comparable calcium entry **(Figure 5G)**, and the increase in oxygen consumption rate was close to significance **(Figure 5H)**. Overall, PKM1 overexpression overcame the metabolic bottleneck and improved glucose-stimulated function in SC-β cells.

**Figure 5.**
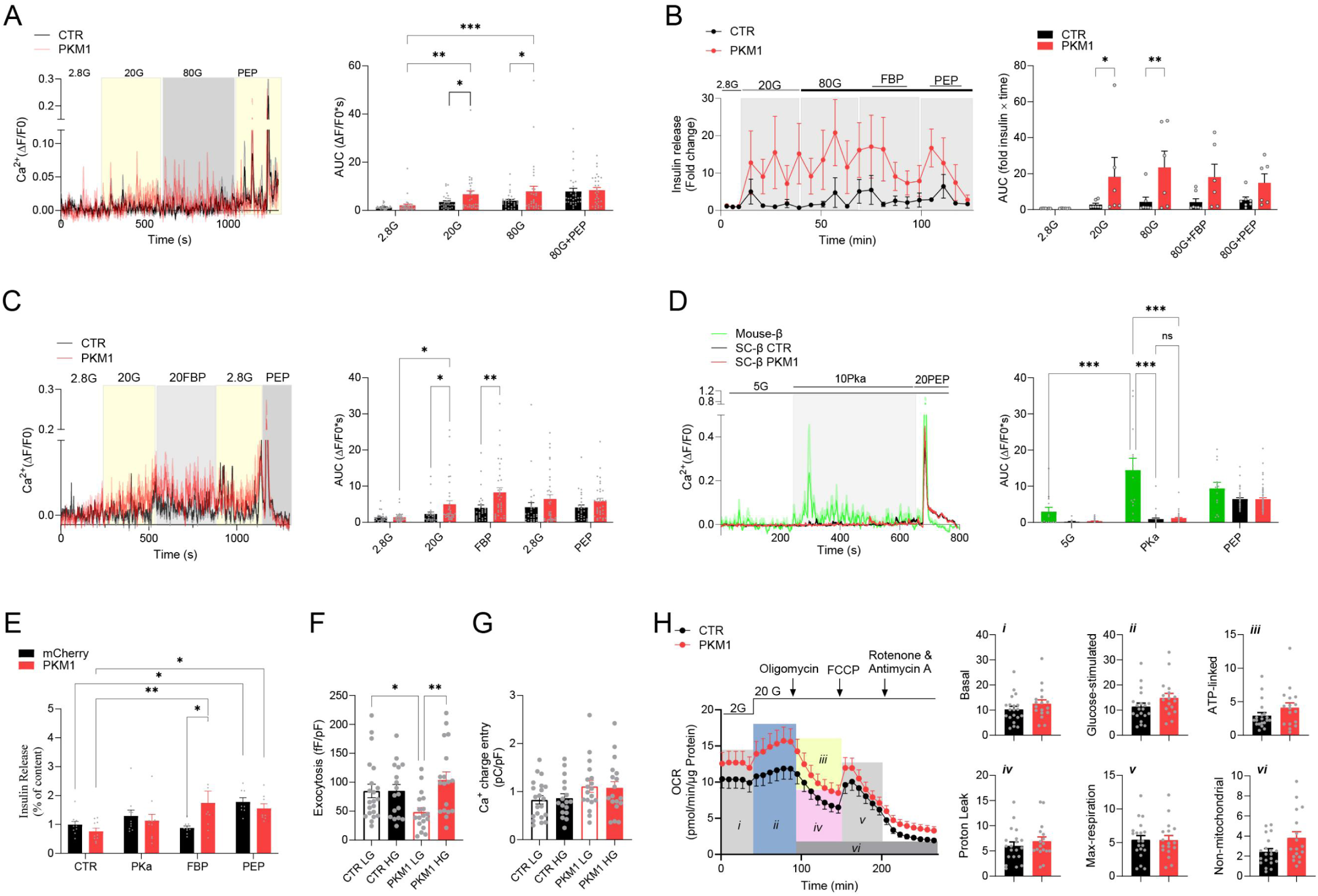
Restoration of *PKM1* expression improves functional response in SC-β cells. **(A)** [Ca^2+^]_i_ recordings to different concentration of glucose level and 20 mM PEP from CTR and *PKM1*-overexpressing SC-β (n=29, 31 cells from 5 different experiments). **(B)** Dynamic insulin secretion to different concentration of glucose, 20 mM FBP and 20 mM PEP from CTR and *PKM1*-overexpressing SC-islets (n=6, 6). **(C)** [Ca^2+^]_i_ recordings to 20 mM FBP and 20 mM PEP from CTR and *PKM1*-overexpressing SC-β (n=29, 34 cells from 4 different experiments). **(D)** [Ca^2+^]_i_ recordings to 10 µM PKa and 20 mM PEP from CTR and *PKM1*-overexpressing SC-β (n=13 mouse-β, n=20 CTR cells, n=20 *PKM1*-overexpressing cells). **(E)** Static insulin secretion to 10 µM PKa, 20 mM FBP and 20 mM PEP at 5 mM glucose from CTR and *PKM1*-overexpressing SC-islets (n=3, 3 independent experiments in triplicate). **(F)** Average total normalized exocytotic responses of CTR and *PKM1*-overexpressing SC-β cell (n=21, 17, 18, 18 cells). **(G)** Average Ca^2+^ currents of CTR and *PKM1*-overexpressing SC-β cell (n=21, 17, 18, 18 cells). LG, 2.8 mM glucose. HG, 20 mM glucose. **(H)** Oxygen consumption rate (OCR) measured by Seahorse assay (n = 6 CTR, 5 *PKM1* in at least triplicate), with relevant respiration parameters calculated at right as shown by italicized numbers. 2G, 2 mM glucose. 2.8G, 2.8 mM glucose. 20G, 20 mM glucose. 80G, 80 mM glucose. FBP, fructose 1,6-bisphosphate. PKa: TEPP-46. CTR: control. SC-β differentiated from H1-Ins-EGFP cells or mouse-β were loaded with Cal-500 AM for calcium imaging in (A,C,D). SC-β differentiated from H1-Ins-EGFP cells was used for patch clamp measurement. *PKM1*-overexpressing SC-β was identified by mCherry overexpression. All data are presented as mean ± SEM, with statistical significance determined using one-way or two-way ANOVA followed by Sidak’s multiple comparison test. *P < 0.05, **P < 0.01, ***P < 0.001.

### PKM1 overexpression promoted SC-**β** cell maturation via regulating PEP metabolism

Since the presence of PEP induced transcriptional changes that hindered SC-β cell maturation, we thereby performed single cell RNA sequencing to explore whether *PKM1* overexpression was sufficient to reverse these transcriptional changes to improve functional maturation. *PKM1*-overexpressing SC-islets were cultured with or without 5 mM PEP throughout Stage 6. These islets were collected at S7D1 for single-cell RNA sequencing (S7D1PKM1-Con and S7D1PKM1-PEP). The resulting data were integrated with single-cell dataset from *PKM1*-overexpressing SC-islets at S7D14 and human adult islets (19,435 cells) (*20*), yielding a combined dataset of 28,168 cells. The cell dataset was clustered according to cell identity. A total of eleven major cell-type classes with marker genes were identified **(Supplementary Figure 3A**-K), including around 25% SC-β cells across SC-islets dataset **(Supplementary Figure 3L)**.

Since *PKM1* cannot be distinguished from *PKM2* in sequencing data, we used mCherry expression as an indicator of successful *PKM1* overexpression. We categorized the dataset into four groups based on *PKM1*-overexpressing SC-islets cultured with or without 5 mM PEP: S7D1PKM1-PEP- (no *PKM1* overexpression, no PEP), S7D1PKM1-PEP+ (no *PKM1* overexpression, with PEP), S7D1PKM1+PEP- (*PKM1* overexpression, no PEP), and S7D1PKM1+PEP+ (*PKM1* overexpression, with PEP) **(Supplementary Figure 4A)**. Consistent with previous findings, PEP treatment downregulated genes and pathways associated with β-cell identity, insulin secretion, and TCA cycle **(Supplementary Figure 4B)**. Notably, many of these downregulated genes and pathways showed a trend toward improvement upon *PKM1* overexpression, suggesting a potential restorative effect of *PKM1* on β-cell function under PEP treatment **(Supplementary Figure 4C)**.

To further explore the transcriptional changes induced by *PKM1*, single-cell sequencing datasets (6,264 cells) generated from *PKM1*-overexpressing islets collected at S7D1 and S7D14 were integrated with adult islets dataset and categorized into four groups based on time of origin and *PKM1*-overexpression: S7D1PKM1-, S7D1PKM1+, S7D14PKM1-, and S7D14PKM1+ **(Figure 6A)**. Compared to S7D1PKM1-, as expected, extended culture of SC-islets to S7D14PKM1-showed a more mature β-cell gene profile (INS, IAPP, HOPX) accompanied by higher gene expression related to insulin secretion (PCSK1, FXYD2, CPE, RBP4, FFAR1, PCSK1N, TTR) (*1*) and OXPHOS pathway (Aerobic respiration and respiratory electron transport) **(Figure 6B, Supplementary Data 4)**, most of which were recapitulated by the comparison between S7D1PKM1+ and S7D14PKM1+ **(Figure 6C, Supplementary Data 4)**. Notably, S7D1PKM1+, compared to S7D1PKM1-, induced similar transcriptional changes and enriched pathways, including aerobic respiration and respiratory electron transport pathway **(Figure 6D, Supplementary Data 4)**, which were further demonstrated in the comparison between S7D14PKM1- and S7D14PKM1+ **(Figure 6E, Supplementary Data 4).** As shown in **(Figure 6F),** genes related to β cell identity, insulin secretion, glycolysis, OXPHOS and exocytosis, such as *INS, PCSK1N, PKM, MT-CO1* and *VAMP2* showed a trend of culture time- and *PKM1*-dependent increase. Consistently, *PKM* expression in adult β-cells, including *PKM1* and *PKM2*, was significantly higher than in non-*PKM1*-overexpressing cells and comparable to the *PKM1*-overexpressing group. Taken together, *PKM1* overexpression triggered a transcriptional program that supports both β-cell functional maturation.

**Figure 6.**
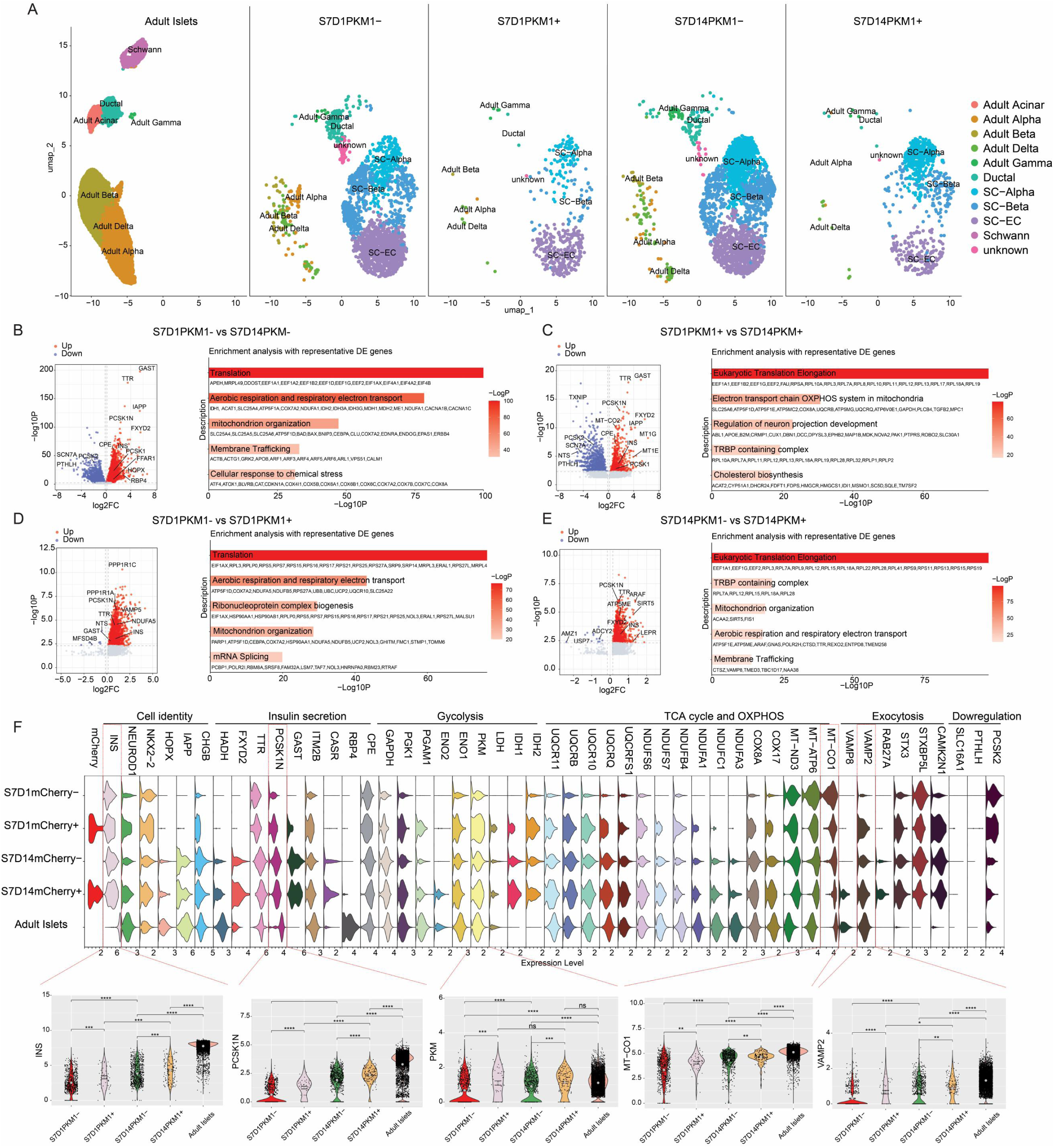
Restoration of *PKM1* expression drives a more functional state in SC-β cells. **(A)** Single-cell RNA-seq transcriptomic profiling of SC-islets overexpressing *PKM1* collected on day 1 of stage 7 (S7D1) and day 14 of stage 7 (S7D14). A UMAP-based embedding projection of an integrated dataset comprising 25,380 SC-derived cells and adult human islet cells, separated by sample origin and clustered by cell type. **(B-E)** Differentially expressed (DE) genes between **(B)** S7D1PKM- and S7D14PKM-**(C)** S7D1PKM+ and S7D14PKM+ **(D)** S7D1PKM- and S7D1PKM+ **(E)** S7D14PKM- and S7D14PKM+ β cell group, including adult and SC-β cells (left). DE genes were submitted to Metascape for enrichment analysis. **(F)** DE signature genes related to cell identity, insulin secretion, glycolysis, TCA cycle and OXPHOS, exocytosis, and those downregulated genes. All data are presented as mean ± SEM, with statistical significance determined using one-way ANOVA followed by Sidak’s multiple comparison test. *P < 0.05, **P < 0.01, ***P < 0.001, ****P < 0.0001.

## Discussion

Aberrant glucose metabolism has been observed in SC-β cells but key metabolic bottleneck that decoupled the glucose response from GSIS remained unknown. Our studies demonstrated that the metabolic bottleneck and immaturity of SC-β cells were caused by elevated PEP at basal glucose, which was linked to the absence of *PKM1*. Additionally, PEP significantly induced calcium responses through a K_ATP_-independent pathway and negatively affected SC-β cell functional state. Restoring *PKM1* promoted the functional maturation of SC-β cells and improved their functional response to glucose.

### Accumulated PEP inhibits GSIS

PEP metabolism is critical for insulin secretion (*9*, *31*). Increasing extracellular glucose from basal to stimulatory levels in primary islets significantly elevated islet PEP content, reaching up to 1 mM (*32*). However, glucose-stimulated increase of PEP was impaired in SC-β cells due to a higher labelling of glycolytic PEP (M+3) at basal glucose. This leads to a loss of glucose-stimulated increase of calcium response and insulin secretion as demonstrated by the following evidence: First, metabolites downstream of PEP did not display glucose-independent increase as those upstream metabolites, which was required for GSIS. Second, exocytosis in SC-β cells was already much higher than in mouse-β cells at basal glucose and cannot be further amplified by 3-PG, 2-PG and PEP. In addition, depolarization-induced calcium entry was higher in SC-β cells. Considering the strong effect of PEP on calcium response at basal glucose, the stimulatory effect by high glucose could be already saturated by elevated PEP. That means the modest glucose-stimulated calcium response and insulin secretion in SC-β cells may not only result from insufficient glycolysis at high glucose levels (*4*), but also from the hyperactivation of calcium response induced by basal PEP. Third, PEP alone induced a transcriptional change that was not favorable for β cell maturation, indicative of a metabolic programming role of PEP.

### PEP metabolism as a metabolic bottleneck of functional maturation

Like previous studies, we used different metabolites to identify the metabolic bottleneck. On one hand, bypassing PEP metabolism with mitochondrial metabolites downstream of PEP, such as membrane-permeable MP and OAA, significantly enhanced insulin secretion (*4*). On the other hand, metabolites upstream of PEP failed to generate insulin secretion or calcium response, indicative of a bottleneck between 2-PG and pyruvate. Lack of *PKM1* led to inadequate pyruvate kinase activity and disrupted the PEP metabolism. A previous study identified a metabolic bottleneck at the GAPDH and PGK1 steps (*4*). However, in our study GAPDH and PGK activity showed only a slight decrease. Additionally, the glucose-dependent increase in 3-PG/2-PG was still maintained, suggesting the presence of an additional glycolytic bottleneck. Similar to PEP metabolism, pyruvate metabolism also exhibited elevated basal pyruvate level and loss of glucose-stimulated increase. Pyruvate cycling is essential in insulin secretion (*25*, *33*, *34*). Recent study reports that pyruvate metabolism is also mediated by LDHB, which is required for inhibiting basal insulin secretion (*35*). However, LDHB level is much lower in SC-islets **(Figure 6F)**, indicative of a pyruvate metabolic bottleneck. In fact, SC-β cells still differ significantly from primary human β cells in the metabolic pathways essential for insulin secretion (*5*). Therefore, identification of a master regulator that upregulates this pathway is crucial for improving functional maturation (*36*).

Surprisingly, elevated basal PEP in SC-β cell raises the basal calcium level more effectively than other metabolites, such as pyruvate or OAA, via a more universal mechanism rather than K_ATP_ channel in SC-β cell. PEP-could induce calcium response possibly through ER, mitochondria or other potassium channels (*37–39*), which warrant further study. In addition to its role in activating calcium, measurements of depolarization-induced exocytosis in mouse β cells supported that PEP, along with 3-PG and 2-PG, is a strong metabolic factor involved in the amplification pathway. Considering the lack of pyruvate kinase activity in SC-β cells, PEP seemed to amplify exocytosis independent of anaplerosis.

One possible mechanism underlying PEP effect on functional maturation could be the impaired Ca^2+^ handling and signaling. Glucose-stimulated calcium influx is not only a hallmark of functional maturation in β cells but also a crucial event in regulating β cell maturation (*15*, *36*). During pancreatic β cell maturation, the calcium influx induced by glucose stimulation precedes the acquisition of insulin secretion ability (*40*). As glucose metabolism matures, proper glucose metabolism triggers calcium influx, activating the calcium-dependent phosphatase Calcineurin (CaN). This activation promotes the dephosphorylation and nuclear translocation of nuclear factor of activated T cells (NFAT), which binds to β cell-specific DNA and regulates transcription of specific genes, promoting insulin vesicle formation and functional maturation in β cells (*41–44*). However, excessive and sustained calcium influx can delink NFAT transcriptional activity, leading to β cell dedifferentiation (*45*). Therefore, calcium mishandling by PEP in SC-β cells may be a key factor impeding functional maturation.

### *PKM1* improves functional state via PEP metabolism

Although *PKM1* overexpression in SC-β cells enhanced glucose-stimulated responses—calcium entry, insulin secretion, oxygen consumption, and exocytosis—this effect should not be solely due to *PKM1*’s enzymatic role in restoring glycolysis-anaplerosis coupling. Given that the transcriptional profile and glucose-stimulus coupling in SC-β cells are still far from those of primary pancreatic β cells, attempts to restore glucose response by acutely fixing one or two metabolic blocks are unlikely to succeed (*4*). We also failed to elicit significant changes of insulin secretion with acute treatment of PKM2 agonist, consistent with previous study (*16*). Additionally, although FBP seemed to enhance insulin secretion in PKM1-overexpressing islets, this effect was likely due to enhanced functional maturation rather than PKM2 activation, as acute exposure to TEPP-46 did not produce a significant effect. We speculate that PKM2 in SC-β cells may be more resistant to activation by FBP or TEPP-46 and considerably less active than in mouse β-cells, as suggested by the predominance of its inactive form in SC-β cells. Furthermore, with minimal *PKM1* expression in SC-β cells, overall pyruvate kinase activity was insufficient to effectively convert PEP to downstream metabolites.

Unlike short-term exposure, chronic TEPP-46 treatment has been shown to enhance functional maturation of SC-β cells (*16*). In fact, long-term *PKM1* overexpression reversed the functional impairment induced by PEP and improved the functional state of SC-β cells by upregulating genetic programs related to cell identity, glycolysis and OXPHOS. Therefore, rather than by just reconnecting glycolysis-anaplerosis coupling, constituent activation of PK activity, either through long-term PKM2 activation or *PKM1* restoration, primarily enhances glucose response by promoting functional maturation via regulating PEP metabolism. Previous studies have individually deleted *PKM1* or *PKM2* to assess their effects on β cell function (*10*). However, these effects appeared to be modest. It would be intriguing to investigate the dual deletion of both *PKM1* and *PKM2* to determine how this impacts PEP levels as well as the functional maturation of SC-β cells.

### Summary

In summary, PEP accumulation disrupts β-cell function by elevating basal calcium levels, thereby suppressing glucose-stimulated insulin secretion (GSIS), decoupling glycolysis from oxidative phosphorylation (OXPHOS), and inhibiting β-cell maturation. *PKM1* overexpression enhances GSIS through a dual mechanism: restoring the glycolysis-OXPHOS linkage and promoting β-cell functional maturation **(Supplementary Figure 5)**. This suggests that *PKM1* plays a crucial role in facilitating β-cell maturation and insulin secretion through metabolic regulation. This study has several limitations. First, due to the scarcity of human islets, mouse islets or β-cells served as positive controls to identify metabolic bottlenecks in SC-islets. Although species-specific differences remain an important consideration, mouse islets offer a reproducible and controlled model system and exhibit many structural and functional similarities to human islets. Second, while *PKM1* overexpression enhances the functional maturation of SC-β cells, the exact mechanisms by which *PKM1* and PEP influence SC-β functional maturation remain unknown. Identifying these mechanisms is essential for advancing our understanding.

---

**Supplementary Figure 1.**
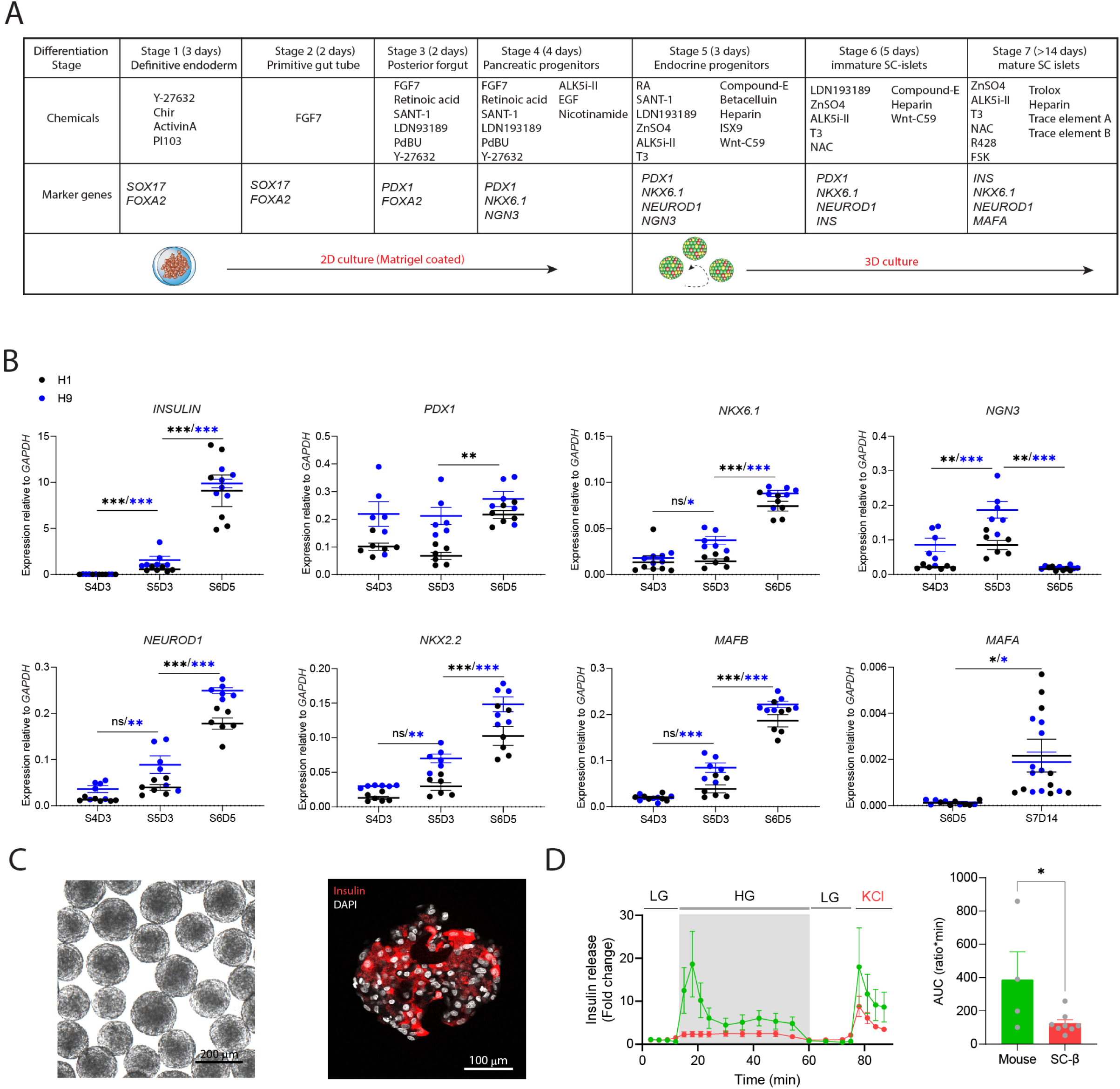
Differentiation and Characterization of SC-islets. **(A)** Overview of 7-stage differentiation protocol to generate SC-islets. **(B)** β cell-specific gene expression of SC-islets differentiated from H1 (n=3 batches of islets) and H9 (n=3, 3 independent experiments in duplicate) cell line at different stages. The color of asterisk and dots correspond to the cell line. **(C)** Cryo-section of SC-islets from S7D14 (left) immunostained for insulin (right). **(D)** Dynamic insulin secretion of mouse- and SC-islets stimulated by LG (low glucose, 2.8 mM), HG (high glucose, 16.7 mM) or KCl (30 mM). mouse islets (n=4) and SC-islets (8) for experiments. All data are presented as mean ± SEM, with statistical significance determined using unpaired Student’s t test or one-way ANOVA followed by Sidak’s multiple comparison test. *P < 0.05, **P < 0.01, ***P < 0.001.

**Supplementary Figure 2.**
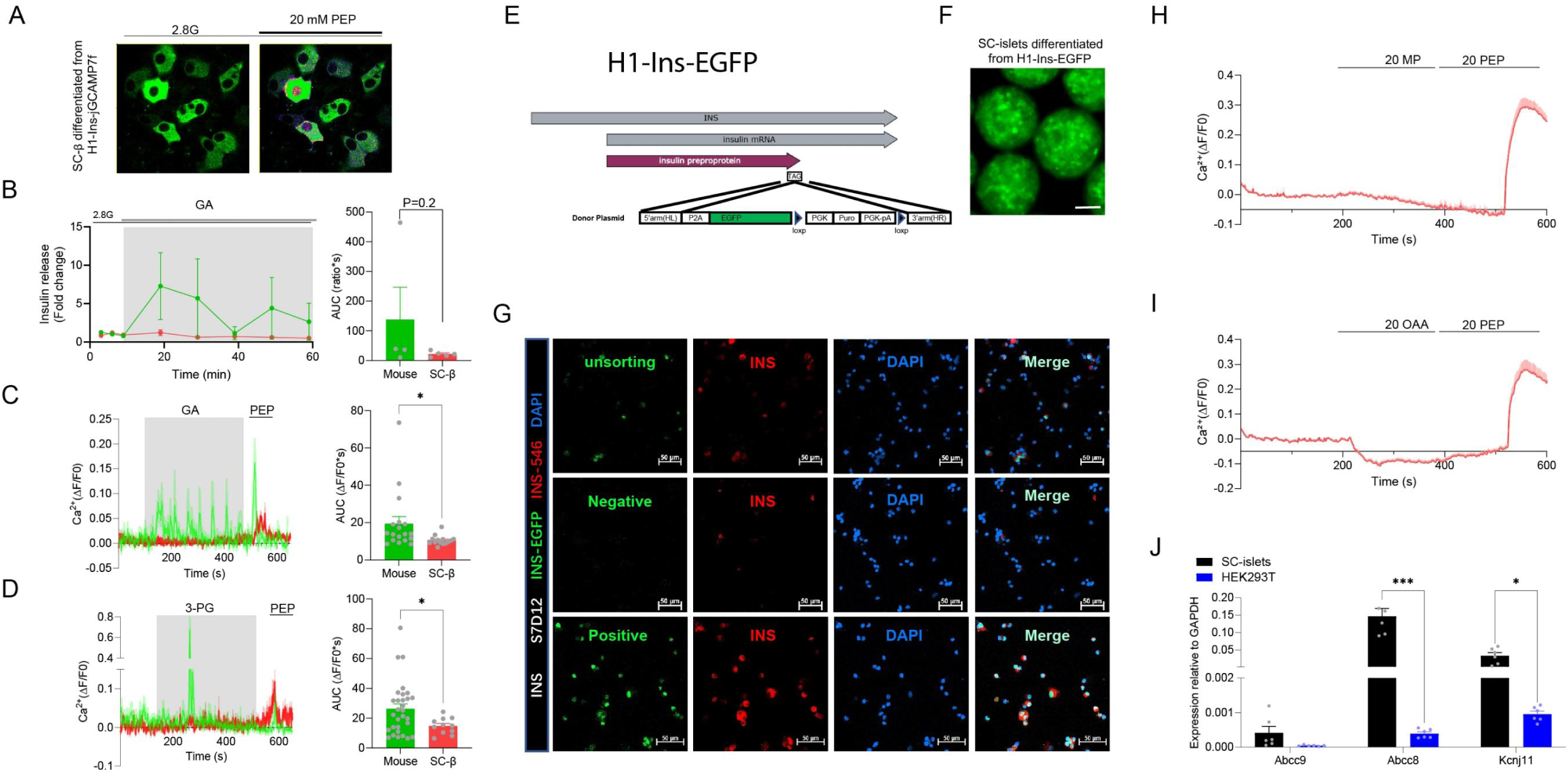
Intermediate metabolite-stimulated functional response. **(A)** Representative [Ca^2+^]_i_ recording of SC-β cell differentiated from H1-Ins-jGCAMP7f to 2.8 mM glucose and 20 mM PEP. Green color indicated the jGCAMP7f Ca^2+^ sensor and fire color indicated the increased fluorescent signal induced by PEP. **(B)** Insulin secretion to 20 mM glyceric acid (GA) at 2.8 mM glucose (2.8G) from mouse-islets (n=4) and SC-islets (n=5). **(C)** [Ca^2+^]_i_ recordings to 20 mM GA followed by 20 mM PEP stimulation from mouse-β (n=18) and SC-β (n=14). **(D)** [Ca^2+^]_i_ recordings to 20 mM 3-PG followed by 20 mM PEP stimulation from mouse-β (n=30) and SC-β (n=11). **(E)** Schematic of generation of H1-Ins-EGFP cell line. **(F)** SC-islets differentiated from H1-Ins-EGFP cells line at S7D14. Scale bar=100 µm. **(G)** Immunostaining of dissociated SC-islet cell for insulin before and after FACS-sorting at S7D12. **(H)** [Ca^2+^]_i_ recordings to 20 mM methyl-pyruvate (MP) followed by 20 mM PEP stimulation from SC-β differentiated from H1-Ins-jGCaMP7f (n=37 cells). **(I)** [Ca^2+^]_i_ recordings to 20 mM oxaloacetic acid (OAA) followed by 20 mM PEP stimulation from SC-β differentiated from H1-Ins-jGCaMP7f (n=37 cells). **(J)** Gene expression level of K_ATP_-related subunits from SC-islets and HEK293T cells (n=3 experiments). All data are presented as mean ± SEM, with statistical significance determined using unpaired Student’s t test or two-way ANOVA followed by Sidak’s multiple comparison test. *P < 0.05, **P < 0.01, ***P < 0.001.

**Supplementary Figure 3.**
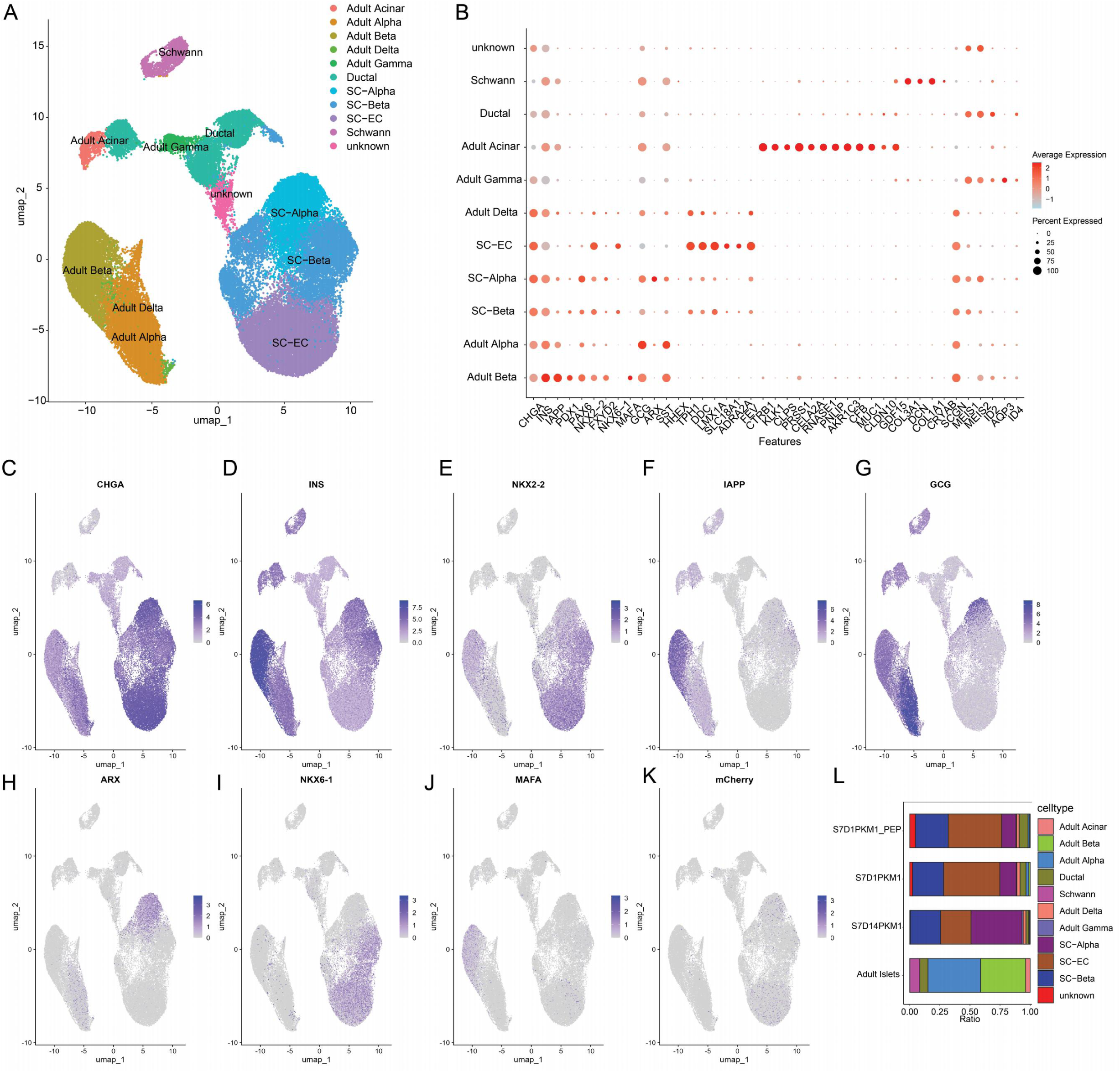
Single-cell RNA-seq analysis of *PKM1*-overexpressing SC-islets PEP-treated with PEP. **(A)** Single-cell RNA-seq transcriptomic profiling of *PKM1*-overexpressing SC-islets treated with PEP collected on day 1 of stage 7 (S7D1) and day 14 of stage 7 (S7D14). A UMAP-based embedding projection of an integrated dataset comprising 25,380 SC-derived cells and adult human islet cells clustered by cell type. **(B)** Signature marker genes for each cell type. **(C-K)** Relative expression of marker genes for endocrine cells (CHGA), α- (GCG, ARX) and β- (INS, NKX2-2, NKX6-1, IAPP, MAFA) cells. mCherry expression indicates the successful expression of *PKM1*. **(L**) Cell number ratio for each cell type in the dataset.

**Supplementary Figure 4.**
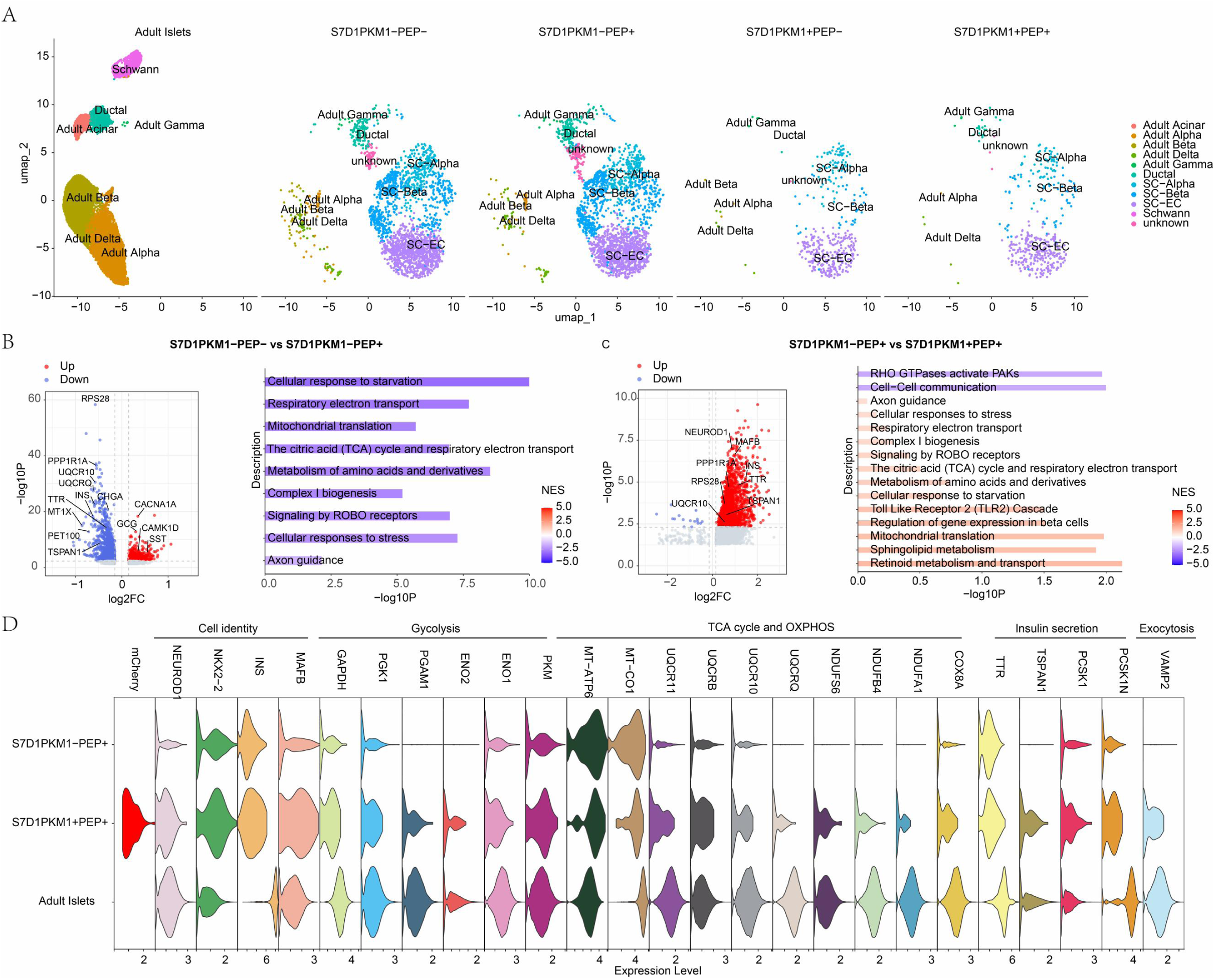
Overexpression of *PKM1* reversed the transcriptional changes induced by PEP. **(A)** Single-cell RNA-seq transcriptomic profiling of *PKM1*-overexpressing SC-islets treated with or without PEP throughout stage 6 and collected on day 1 of stage 7 (S7D1). A UMAP-based embedding projection of an integrated dataset comprising 5,525 SC-derived cells 19,435 adult human islet cells, separated by sample origin and clustered by cell type. **(B)** DE gene comparison of S7D1PKM1-PEP- and S7D1PKM1-PEP+, which integrates the dataset of S7D1Con and S7D1PEP from (figure 3J) and the dataset of S7D1PKM and S7D1PKM1_PEP to make a larger dataset (left). Gene sets enriched analysis (GSEA) using Reactome pathways showed upregulated and downregulated pathways along with DE genes. **(C)** DE gene comparison of S7D1PKM1-PEP+ and S7D1PKM1+PEP+ from the dataset of S7D1PKM-Con and S7D1PKM1-PEP (left). Gene sets enriched analysis (GSEA) using Reactome pathways showed upregulated and downregulated pathways along with DE genes between. NES, normalized enrichment score. DE, differentially expressed. **(D)** DE signature genes between S7D1PKM1-PEP+ and S7D1PKM1+PEP+ related to cell identity, insulin secretion, glycolysis, TCA cycle and OXPHOS, exocytosis. All data are presented as mean ± SEM.

**Supplementary Figure 5.**
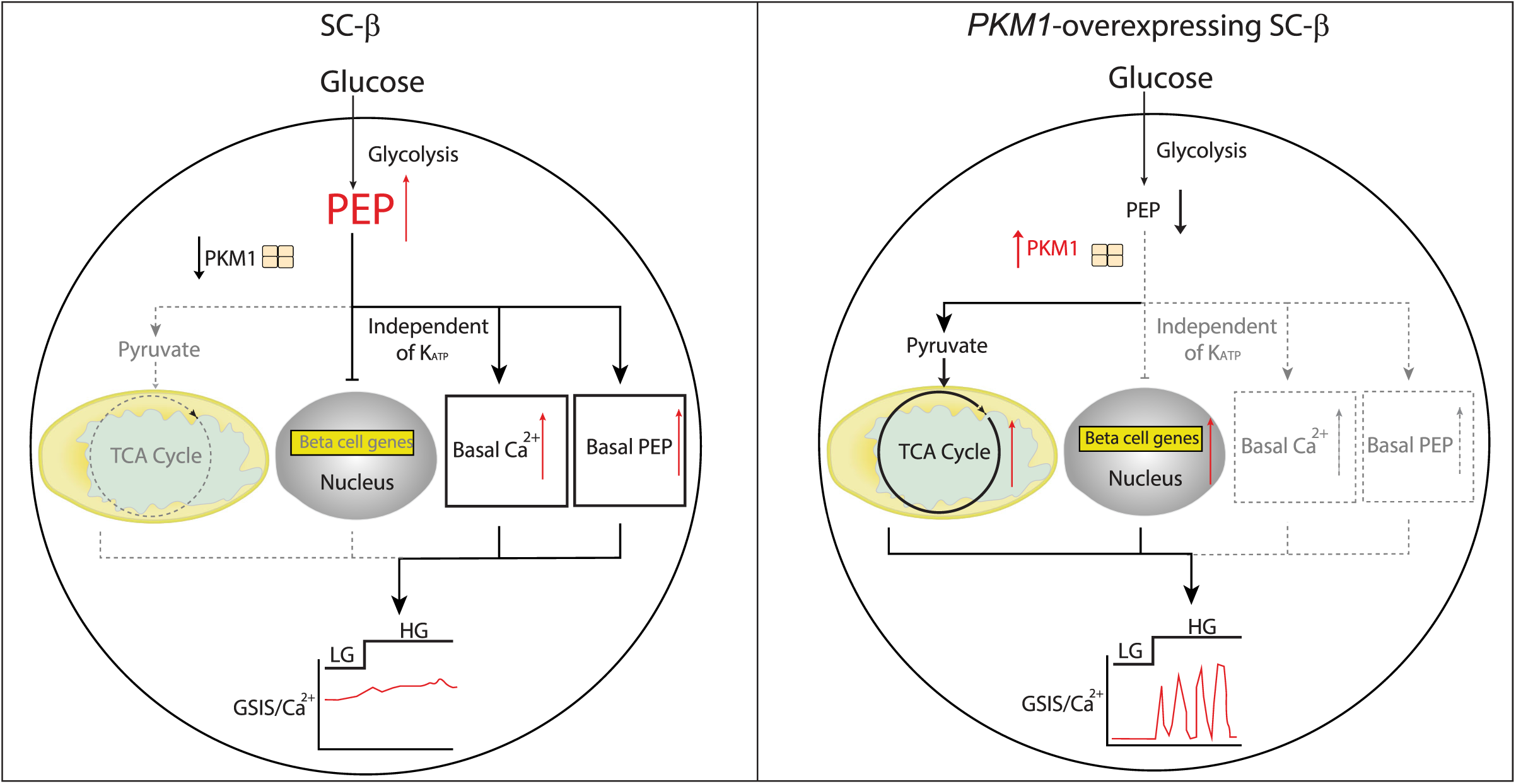
Schematic depicting the role of PEP accumulation and PKM1 overexpression in SC-β cell functional maturation. Immature SC-β cells display constituently high level of basal PEP due to the low expression of PKM1. Elevated basal PEP prevents glucose-stimulated changes of downstream metabolites, which are further aggravated by the absence of PKM1 and lead to weak glucose-stimulated calcium response and insulin secretion. Furthermore, PEP alone raises basal calcium level independent KATP and blocks functional maturation. Overexpression of PKM1 is sufficient to reverse the metabolic programming induced by PEP to improve functional maturation and glucose-stimulated function.

## Acknowledgments

we would like to thank the Proteomics and Metabolomics Platform, Guangzhou Laboratory for our liquid chromatography-time-of-flight mass spectrometry (Q-TOF) and we would be grateful to Nannan Wang for her help of data collection and analysis.

## Author contributions

Huisheng Liu, Zonghong Li, Yanying Guo and Tao Xu conceived and supervised the study. Haopeng Lin, Deqi Chen, Feng Zhang, Xin Liu, Wei Peng, Lihua Chen designed and performed experiments, data analysis and interpretation. Kuo Jiang and Peng Xue performed single cell transcriptomics analyses and interpretation. Qifei Dong and Jiawei Yan participated in experiments related to stem differentiation, gene expression measurement and calcium imaging. Jiaxiang Yin and Zirong Bi designed the perifusion apparatus for dynamic insulin secretion and calcium imaging. Feng Zhang, Deqi Chen, Xin Liu and Tongran Zhang established the reporter cell lines. Deqi chen, Xin Liu, Xiaoxiao Xie participated in mouse management and islet isolation. Huisheng Liu and Haopeng Lin wrote the paper, with contributions from all authors.

## Data and materials availability

All data are available in the main text or the supplementary materials.

## Materials and Methods

### Cell culture and differentiation of SC-β cells

#### In vitro differentiation of H1 and H9

H1 or H9 **human embryonic stem cells (hESC)** were cultured on 1:100 diluted Matrigel (Corning, 354277) in mTeSR medium (Stemcell, 85850). After rinsing with Mg^2+^/Ca^2+^-free DPBS (Thermo Fisher, 14190250) and digestion with TrypLE Express Enzyme (1×) (Thermo Fisher, 12605028) for 3 minutes at 37°C, the digestion was stopped with mTeSR. Single cells were collected and spun at 1,000 rpm for 5 minutes. The cell pellet was resuspended in mTeSR medium supplemented with Y-27632 (10 mM; MCE, HY-10583), and single cells were seeded at approximately 1 × 10^5^ cells/well on Matrigel-coated surfaces. Cells were fed with mTeSR medium daily and differentiated after 48 hours at approximately 90% confluency. The differentiation protocol was modified based on previous protocols (*2*, *17*) (see also **Supplementary Data 1**).

##### S1

Definitive endoderm (3 d). Undifferentiated pluripotent stem cells plated on 1:100 Matrigel-coated surfaces were cultured in MCDB 131 medium (Reprocell, PM151210) with 0.5% BSA (fatty acid-free BSA, Proliant, 69760), 1.7 g/l sodium bicarbonate (SIAL, S6014), 10 mM final glucose concentration (Sigma, G7528), 0.25 mM Vitamin C (Sigma, A1300000), 1% Glutamax (Thermo Fisher Scientific, 35050061), 1% Penicillin-Streptomycin (Thermo Fisher Scientific, 15140122), 100 ng/ml Activin A (MCE, HY-P70311), 3 μM ChiR99021 (MCE, HY-10182), 50 nM PI-103 (Selleck, S1038), and 10 μM Y-27632 for day 1 only. For day 2, cells were cultured in MCDB131 with 0.5% BSA, 1.7 g/l sodium bicarbonate, 10 mM glucose, 0.25 mM Vitamin C, 1% Glutamax, 1% PS, 50 ng/ml Activin A, and 0.1 μM ChiR99021. On day three, cells were cultured in MCDB with 0.5% BSA, 1.7 g/l sodium bicarbonate, 10 mM glucose, 0.25 mM Vitamin C, 1% Glutamax, 1% PS, and 50 ng/ml Activin A.

##### S2

Primitive gut tube (2 d). S1 cells were transferred to MCDB 131 medium containing 0.5% BSA, 1.7 g/l sodium bicarbonate, 10 mM glucose, 0.25 mM Vitamin C, 1% Glutamax, 1% PS, and 50 ng/ml FGF7 (Stemcell, 78046.2) for 2 days.

##### S3

Posterior foregut (2 d). S2 cells were cultured for an additional 2 days in MCDB 131 medium supplemented with 2% BSA, 2.5 g/l sodium bicarbonate, 10 mM glucose, 0.25 mM Vitamin C, 1% Glutamax, 1% PS, 1:200 ITS-X (Reprocell, pb180431), 50 ng/ml FGF7, 2 μM retinoic acid (Sigma, R2625), 0.25 μM SANT-1 (MCE, HY-100224), 100 nM LDN193189 (BMP receptor inhibitor, MCE, HY-12071), 500 nM PdBu (Millipore, 524390), and 10 μM Y-27632.

##### S4

Pancreatic endoderm (4 d). S3 cells were further differentiated in MCDB 131 medium supplemented with 2% BSA, 2.5 g/l sodium bicarbonate, 10 mM glucose, 0.25 mM Vitamin C, 1% Glutamax, 1% PS, 1:200 ITS-X, 2 ng/ml FGF7, 2 μM RA, 0.25 μM SANT-1, 100 nM LDN193189, 100 nM PdBu, 10 μM Y27632, 10 μM ALK5 inhibitor II (Selleck, s7223), 100 ng/ml EGF (MCE, HY-P7109), and 10 mM Nicotinamide (Sigma-Aldrich, N0636) for 4 days.

##### S5

pancreatic endocrine precursors (3 d). On the last day of S4 culture, cells were treated with accutase (Stemcell, 7920) for 8 min at 37 °C, followed by gentle pipetting to break into single cell suspension. Next, cell suspension were transferred into low-attached 6-well culture plate in suspension at 100 rpm with S5 medium, which consisted of MCDB 131 medium with 2% BSA, 2 g/l sodium bicarbonate, 20.6 mM glucose, 0.25 mM Vitamin C, 1% Glutamax, 1% PS, 1:200 ITS-X, 0.05 μM RA, 0.25 μM SANT-1, 100 nM LDN193189, 10 μM zinc sulfate (ZnSO_4_, Sigma, Z0251), 10 μM ALK5 inhibitor II, 1 μM T3 (MCE, HY-A0070), 1 μM compound E(MCE, HY-14176), 20 ng/ml Betacellulin (Stemcell, 78105), 10 μg/ml Heparin (Sigma, H3149), 10 μM Isoxazole 9 (ISX9, MCE, HY-12323) and 100 nM Wnt-C59 (MCE, 1243243-89-1) for 3 dads. For lentivirus infection, mcherry or *PKM1*-mCherry lentivirus were added at S5D1, achieving stable *PKM1* expression at S5D3. After 48h, lentivirus media was replaced with new S5 media.

##### S6

S6 cells, characterized by NKX6.1 and insulin expression, were cultured for 5 days. The culture medium consisted of MCDB 131 supplemented with 2% BSA, 1.8 g/l sodium bicarbonate, 20.6 mM glucose, 0.25 mM Vitamin C, 1% Glutamax, 1% PS, 1:200 ITS-X, 100 nM LDN193189, 10 μM ZnSO4, 10 μM ALK5 inhibitor II, 1 μM T3, 0.1 μM compound E, 10 μg/ml Heparin, 100 nM Wnt-C59, and 1 mM N-acetylcysteine (Sigma, A9165) for 5 days. For PEP treatment, 5 mM PEP was added to media and media pH was corrected with NaHCO_3_ as that of S6 media.

##### S7

S7 cells, characterized by NKX6.1, insulin, and MAFA expression, were cultured for 14-21 days. The culture medium consisted of MCDB 131 supplemented with 2% BSA, 1.5 g/l sodium bicarbonate, 5.6 mM glucose, 1% Glutamax, 1% PS, 1:200 ITS-X, 10 μM ZnSO4, 10 μM ALK5 inhibitor II, 1 μM T3, 10 μg/ml Heparin, 1 mM N-ace, 10 μM Trolox (Sigma, T3251), 1:2000 Trace element A (Corning, 25-021-CI), 1:2000 Trace element B (Corning, 25-022-CI), 2 μM R428 (SelleckChem, S2841), 1:2000 chemically defined lipid concentrate (lipid, Gibco, 11905031), and 10 μM Forskolin (Aladdin, 66575-29-9).

### Generation of reporter cell line

H1-Ins-EGFP wase generated as previously reported for H1-Ins-jGCaMP7f (Liu et al., manuscript in submission to Stem Cell Research). We utilized low off-target CRISPR/Cas9n technology for in situ insertion of the EGFP gene in human embryonic stem cells (H1). By employing the Cas9 D10A nickase (PX335, Addgene #42335) with dual sgRNAs targeting each side of the insulin gene stop codon, we inserted P2A-jGCaMP7f-PGK-Puro-pA. This insertion enables simultaneous insulin protein expression for identifying insulin-producing cells in differentiated pancreatic endocrine cells. Cells were transfected at 70% confluence using Lipofectamine™ Stem Reagent (STEM00008), with puromycin added after 24 hours at 1 µg/ml to select for successful insertions, ensuring elimination of non-transfected cells.

### Generation of lentivirus

All overexpression lentiviral plasmids were constructed via homologous recombination. During the construction process, the following steps were undertaken: 1. puc57 was consistently used as the plasmid backbone; 2. The lentiviral integration site from the lentiCRISPR v2 plasmid (addgene: #52961) was utilized; 3. The EF1a promoter was employed to enhance gene expression, followed by the target gene’s ORF (with the stop codon removed), linked to P2A and mCherry for monitoring plasmid transfection efficiency and viral overexpression in 293T cells; 4. All plasmids were recombined using the EZ-HiFi Seamless Cloning Kit (Genstar #T196), and the accuracy of the sequences was verified by DNA sequencing. Pspax2, VSVG, and PLV were mixed in a ratio of 1:0.5:1 for transfection of 293T cells using PEI. 12 hours post-transfection, the medium was replaced with DMEM containing 5% FBS and antibiotics. At 48- and 72-hour post-transfection, the supernatants were collected, centrifuged at 2000 rcf for 10 minutes to remove cell debris, sterile filtered through a Millex 0.22 µm filter, and the viruses were finally concentrated using an Amicon® Ultra filter (#UFC9100).

### Immunohistochemistry

For cryo-sectioning, S7 organoids were fixed in 4% PFA (Sangon Biotech) at room temperature for 1 hour, rinsed with PBS, and incubated overnight at 4°C in 30% sucrose. They were then embedded in OCT, snap-frozen in liquid nitrogen, and stored at −80°C. Sections were cut at 10 μm thickness and mounted on Superfrost Plus slides. For immunostaining of re-plated cells, SC-islets were dissociated, plated on Matrigel-coated confocal dishes and fixed with 4% PFA at 4D overnight. Cells were then permeabilized with Quickblock™ Blocking Buffer (BEYOTIME) at room temperature for 1 hour. Primary antibodies were incubated overnight at 4D, followed by PBS washes, and staining with secondary antibodies and Hoechst 33342 (Thermo Fisher Scientific) for 2 hours at room temperature. Imaging was done on a NIKON A1 confocal microscope. Antibody information is listed in **Supplementary Data 6**.

### Insulin secretion

For dynamic insulin secretion, 25 (mouse islets) or (SC-islets) were handpicked into home-made organoid chip for dynamic insulin secretion as previously described (*47*). Islets were pre-perifused for 1 h at KRB solution with 2 mM or 2.8 mM glucose followed by stimulation of different levels of glucose, permeable metabolites and drugs at a sample collection rate of 10 ml/min every 3-6 minutes. For static insulin secretion, 15 islets per triplicates were preincubated with 2.8 mM for 1 h followed by stimulation of indicated conditions for 1 h. The pH of all solutions containing metabolites or drugs was adjusted to 7.4 using NaOH. D-mannitol (Aladdin Scientific, Shanghai, China) was used to balance the osmolarity of KRB solutions as previously reported (*48*). The samples and lysates of islets were stored at −80 °C until further assayed by Insulin Detection Kit (STELLUX® Chemi Rodent Insulin ELISA kit; STELLUX® Chemi Human Insulin ELISA kit, Alpco). Information of metabolites and drug is listed in **Supplementary Data 7**.

### Intracellular Ca^2+^ and membrane potential imaging

For Ca^2+^ imaging, islets or dissociated cells cultured on 35 mm glass bottom were loaded with Cal-500AM (2.5 μM, AAT Bioquest, USA) or Fluo-8AM (2.5 μM, AAT Bioquest, USA) for 30 min and perifused with pH-corrected and osmolarity-balanced KRB solution at indicated conditions. The fluorescence signal was excited and detected at an excitation/emission wavelength of 405 nm/500 nm for Cal-500AM, and at 490 nm/514 nm for both Fluo-8AM and H1-jGCAMP7f cells. For membrane potential imaging, islets or dissociated cells were loaded with FluoVolt (Thermo Fisher, USA) with an excitation/emission wavelength of 490 nm/514 nm. Ca^2+^ and membrane potential were imaged at 1 Hz on an inverted confocal microscope (Carl Zeiss LSM 980) for single cells and an upright confocal microscope (Carl Zeiss LSM 980 NLO) for whole islets. Mouse β-cells were marked and identified by immunostaining and SC-β cells were identified by using H1-Ins-jGCAMP7f or H1-Ins-EGFP for experiment. Fluorescence ratios were calculated using a tool box from Matlab with indicated analytical method (average fluorescence and smooth dynamics) (*46*).

### ^13^C-glucose labeling metabolomics

∼400 primary mouse islets or SC-islets were used for each replicate and preincubated at KRB solution with 2.8 mM glucose for 60 min, followed by incubation at 2.8 (low) or 16.7 mM (high) [U-¹³CD] glucose (Cambridge Isotope Laboratories, CLM 1396) for 60 min at 37 °C and 5% CO_2_. Afterwards, islets were washed with PBS 3 times and added 50 ul “extraction solution” (Acetonitrile/Methanol/Aqueous=4:4:2 kept at −20’C overnight before using. The mixture was kept in ice to sonicate for 5min followed by future centrifugation at 12000 g for 5min. Supernatant was transferred to vials and run with 4 ul injection volume by LC-MSMS system. Chromatographic separation was performed using an Agilent 1290II ultra-high-pressure liquid chromatography (UHPLC) system equipped with 6546 quadrupole time-of-flight (QTOF) mass spectrometry. A Waters ACQUITY UPLC BEH Amide column (2.1 mm × 100 mm × 1.7 μm) and guard column (2.1 mm × 5 mm × 1.7 μm) at 35 °C was used to separate metabolites with mobile phase A: 100% aqueous containing 15mM ammonium acetate and 0.3% ammonium hydroxy and mobile phase B: 90% acetonitrile (v/v) aqueous containing 15mM ammonium acetate and 0.3% ammonium hydroxy. The linear gradient was set as follows: 10% A (0.0– 8.0 min), 50% A (8.0–10.0 min), 50% A (10.0–11.0 min) and 10% A (11.0–20.0 min). The total run time was 20 min and flow rate was 0.3 mL/min. The mass spectrometer was equipped with Agilent Jet-stream source operating in negative and positive ion mode with source parameters set as follow: Nebulizer gas, 45psi; Sheath gas temperature, 325℃; Sheath gas flow, 10L/min; Dry gas temperature, 280℃; Dry gas flow, 8L/min; Capillary voltage, 3500 V for two ion modes and nozzle voltage, 500 V for positive and 1000 V for negative mode. The QTOF scan parameters were set as follows: Scan speed, 1.5 scan/s; scan range, 50-1700 m/z and ion fragmentor voltage, 140v. The acquired data quality was monitored by Amino Acid standards (Merck, Sigma-Aldrich Production GmbH, Switzerland), mix-samples quality controls and blanks. Peak integration and metabolite isotopologue identification was accomplished using Profinder 10.0 (Agilent). Nature abundance was assayed using non-labelled samples and removed to avoid any possible confounding effect.

### Patch Clamp Analysis

Patch Clamp experiment was performed as previously reported (*49*). Briefly, dispersed single cells cultured on 35 mm dishes overnight were preincubated at 2.8 mM glucose for 1 hour and patched in bath solution containing (in mM): 118 NaCl, 5.6 KCl, 20 TEA, 1.2 MgCl_2_, 2.6 CaCl_2_, 5 HEPES at indicated glucose level with a pH of 7.4 (adjusted by NaOH) at 32-35°C. Whole-cell capacitance was recorded with the sine+DC lock-in function of an EPC10 amplifier and PatchMaster software (HEKA Electronics). Exocytotic responses and inward Ca^2+^ currents were measured 1-2 minutes after obtaining the whole-cell configuration in response to ten 500 ms depolarizations to 0 mV from a holding potential of −70 mV. Changes in capacitance and integrated Ca^2+^ charge entry were normalized to cell size (fF/pF and pC/pF, respectively). The intracellular solution contained (in mM): 125 Cs-Glutamate, 10 CsCl, 10 NaCl, 1 MgCl2, 5 HEPES, 0.05 EGTA, 3 MgATP and 0.1 cAMP with pH=7.15 (pH adjusted with CsOH). For dialysis/infusion experiment, metabolites were added to pipette solution (pH-corrected) as indicated. Measurement was performed 3 minutes after whole cell mode, which achieved homogenous distribution of metabolites. SC-β cells were identified by EGFP fluorescent signal. Mouse β-cells were identified by size (>4 pF) and a Na^+^ channel half-maximal inactivation at around - 90 mV.

### Enzymatic activity measurement

Pyruvate kinase activity was measured using the CheKine™ Micro Pyruvate Kinase (PK) Assay Kit (KTB1120) according to the manual. PK catalyzes phosphoenolpyruvate and ADP reaction into ATP and pyruvate, and lactate dehydrogenase further catalyzes the production of lactic acid and NAD + by NADH and pyruvate. The rate of NADH decline at 340 nm can reflect PK activity. Cell lysates from 400 islets were sonicated on ice using the provided Extraction Buffer and quantified by BCA assay. 10 μL sample, 10 μL Working Reagent and 180 μL Substrate Mix Working Reagent were mixed and added to the 96-well UV plate. Measure the absorbance value at 340 nm with a microplate reader. The data shown represent the first ten data points of NADH consumption in this reaction.

PGK1 activity was measured using the Grace Biotech Micro PGK Assay Kit (G0885W) according to the manual. Phosphoglycerate kinase (PGK) catalyzes the reaction between 3-phosphoglycerate and ATP to produce 1,3-bisphosphoglycerate and ADP. In a subsequent reaction, 1,3-bisphosphoglycerate is catalyzed by glyceraldehyde-3-phosphate dehydrogenase (GAPDH) and reacts with NADH to generate 3-phosphoglyceraldehyde, NADD, and inorganic phosphate. Since NADH absorbs light at 340 nm, the decrease in NADH concentration during the reaction results in a reduction in absorbance at this wavelength. This absorbance change at 340 nm can be used to monitor PGK activity. Cell lysates from 400 islets were sonicated on ice using the provided Extraction Buffer and quantified by BCA assay. 20 μL sample was mixed with the Working Reagent added to the 96-well UV plate. After 10 minutes, measure the absorbance value at 340 nm with a microplate reader. The data shown represent the first ten data points of NADH consumption in this reaction.

GAPDH activity was determined using the Abcam GAPDH colorimetric assay kit (ab204732) as previously reported (*4*). This assay tracks the conversion of the substrate glyceraldehyde-3-phosphate to 1,3-bisphosphoglycerate along with NADH production. NADH, generated by GAPDH activity present in cell lysates, is linked to a proprietary developer included in the kit, which utilizes the NADH from GAPDH to initiate a secondary colorimetric reaction in each well. Cell lysates from 400 islets were sonicated on ice using the provided Extraction Buffer and quantified by BCA assay. After adding 50 µL sample into the reaction mix, the plate was incubated at 37°C for the assay duration. NADH accumulation was indirectly measured at 450 nm by monitoring the developer reaction product in kinetic mode over a 10-minute period. The data presented display the raw conversion values for the first 10 time points of the reaction.

### Oxygen consumption

The Agilent Seahorse XFE24 Analyzer with islet capture microplates was used, following a protocol previously outlined (*22*). Specifically, 70 islets were loaded per well and treated sequentially with 2 mM glucose, 20 mM glucose, 5 µM oligomycin, 5 µM FCCP, and 5 µM rotenone/antimycin A. For each data point, the experiment included a 3-minute mixing phase, a 2-minute waiting phase, and a 3-minute measurement phase. Each biological sample had 3∼4 technical replicates, and the oxygen consumption rate was normalized islet protein content.

### scRNA sequencing analysis

scRNA sequencing analysis were performed as previously reported (*50*). Briefly, approximately 10,000 cells were captured, reverse-transcribed, barcoded, and amplified. cDNA libraries were quantified with Qubit 3.0, analyzed on Agilent 2100, and sequenced on the Illumina NovaSeq 6000 (PE150) by Annoroad Gene Technology. Raw FASTQ files were processed with Cell Ranger (v4.0.0) (https://support.10xgenomics.com/single-cell-gene-expression/software/pipelines/latest/advanced/references) using default settings, aligning reads to the GRCh38 human genome.We included 12 samples of primary adult human pancreatic islets as reference scRNAseq dataset for further analysis (*20*). High quality cells (>300 genes, <10% mitochondrial genes, <3%red blood cell gene expression) were selected for data normalization, scaling and principal component analysis (PCA) with default methods. Top 3000 variable genes were identified from each sample. The Harmony method was applied on the first 30 principal components, using sample as a covariate to integrate different samples (*51*). The integrated principal components were used to construct the UMAP, identify neighboring cells (via shared nearest neighbor), and define cell clusters using default Seurat methods. The clustering resolution was adjusted to 0.5. Differentially expressed genes among clusters were identified using Seurat’s FindMarkers function. The annotation of the clusters was performed using the GPTCelltype package according to the known marker genes (*52*). Adult and SC-β cells were used for identifying differentially expressed gene with a P value < 0.01 and log2 fold change >0.2. We performed gene set enrichment analysis using Reactome (*53*) and Metascape v.3.5 (*54*) with a P value cutoff of <0.05. Detailed analytical result related to differentially genes and enrichment can be seen in **supplementary data 2-4**.

### Fluorescence activated cell sorting

For SC-β (H1-Ins-EGFP) cell sorting, cell aggregates were dissociated into a single-cell suspension using Accutase and resuspended in S7 media containing 10 μM Y27632. EGFP-positive cells were sorted into 15 ml Falcon tubes containing S7 medium and 10 μM Y27632 using a tube sorting mode (MA900, Sony).

### Quantitative RT-PCR

RNA was extracted with the RNeasy Mini Kit (Qiagen) and treated with DNase. cDNA was synthesized using the RT reagent Kit with gDNA Eraser (TAKARA). Real-time qPCR was performed with SYBR Premix Ex TaqII (TAKARA) and analyzed using the ΔΔCt method, normalizing expression to GAPDH. Primer details are in **Supplementary Data 5**.

### Western blotting

SC-islets from S7D14 and mouse islets were collected in RIPA lysis buffer with protease inhibitor (Beyotime). Lysates were centrifuged, and supernatants were separated by SDS-PAGE, transferred to PVDF membranes, blocked, and incubated with primary and HRP-conjugated secondary antibodies. Detection was performed using ECL reagent (Millipore) and captured on a G-Box. Protein quantification was done with ImageJ, normalizing target protein intensity to housekeeping proteins (beta-actin). Antibodies are detailed in **Supplementary Data 6**.

### Data availability

Raw data and count matrices of scRNA-seq data are available under the accession number GSE281603. All other data are available from the corresponding authors upon request. Source data are provided with this paper.

### Code availability

The code for analyzing 10X Genomics sequencing data is available at Seurat (https://satijalab.org/seurat/). Additional detailed code for specific data analysis steps can be provided upon request.

### Analysis and statistics

Data analysis was conducted using FitMaster (HEKA Electronik) and GraphPad Prism (v10.0c). Statistical outliers were identified and removed using the unbiased ROUT (robust regression and outlier identification) test. For normally distributed data, statistical comparisons were made using a 2-tailed Student’s t-test (for two groups) or ANOVA followed by Sidak’s multiple comparison test, as specified in the figure legends.

